# Quantitative trait and transcriptome analysis of genetic complexity underpinning cardiac interatrial septation in mice using an advanced intercross line

**DOI:** 10.1101/2022.10.31.514499

**Authors:** Mahdi Moradi Marjaneh, Edwin P. Kirk, Ralph Patrick, Dimuthu Alankerage, David T. Humphreys, Gonzalo Del Monte-Nieto, Paola Cornejo-Paramo, Vaibhao Janbandhu, Tram B. Doan, Sally L. Dunwoodie, Emily S. Wong, Chris Moran, Ian C.A. Martin, Peter C. Thomson, Richard P. Harvey

## Abstract

Unlike single-gene mutations leading to Mendelian conditions, common human diseases are likely emergent phenomena arising from multilayer, multiscale and highly interconnected interactions. Atrial and ventricular septal defects are the most common forms of cardiac congenital anomalies in humans. Atrial septal defects (ASD) show an open communication between left and right atria postnatally, potentially resulting in serious hemodynamic consequences if untreated. A milder form of atrial septal defect, patent *foramen ovale* (PFO), exists in about one quarter of the human population, strongly associated with ischaemic stroke and migraine. The anatomic liabilities and genetic and molecular basis of atrial septal defects remain unclear. Here, we advance our previous analysis of atrial septal variation through quantitative trait locus (QTL) mapping of an advanced intercross line (AIL) established between the inbred QSi5 and 129T2/SvEms mouse strains, that show extremes of septal phenotypes. Analysis resolved 37 unique septal QTL with high overlap between QTL for distinct septal traits. Whole genome sequencing of parental strains identified high confidence candidate deleterious variants, including in known human congenital heart disease genes, whereas transcriptome analysis of developing septa revealed networks involving ribosome, nucleosome, mitochondrial and extracellular matrix biosynthesis underlying septal variation. Analysis of variant architecture across different gene features, including enhancers and promoters, provided evidence for involvement of non-coding as well as protein coding variants. Our study provides the first high resolution picture of genetic complexity and network liability underlying common congenital heart disease, with relevance to human ASD and PFO.

## Introduction

Congenital heart disease (CHD) is common with a neonatal incidence of 0.8-1%, and places an enormous burden on affected patients, families, and healthcare systems (van der Linde et al., 2011). High throughput sequencing has identified abundant examples of monogenic syndromic and non-syndromic forms of CHD; however, the majority represent complex multifactorial conditions of unknown etiology, with a significant role for polygenic disease, *de novo* mutations and gene-environment interactions (Sifrim et al., 2016). Genome-wide association studies (GWAS) have identified a small number of common CHD risk alleles, likely those of the largest effect (Lahm et al., 2021); however, for the majority of clinical subcategories none have been identified thus far.

Development of the four-chambered mammalian heart leads to permanent separation of the systemic and pulmonary circulations through septation of common atrial and ventricular chambers. Septation co-evolved with air breathing and involves convergence of myogenic and non-myogenic tissues at the valvuloseptal apparatus of the atrioventricular (AV) junction (Moorman and Christoffels, 2003). This process is genetically vulnerable as septation defects occur commonly in the human CHD spectrum (Gruber and Epstein, 2004).

During fetal life, the inter-atrial septum functions initially as a one-way “flap valve” to help divert systemic blood away from the pulmonary circulation (**Figure 1A**). Regulatory networks generated at the boundaries between different cardiac progenitor fields define the programs for septal development (De Bono et al., 2018; Rana et al., 2014; Steimle et al., 2018). Initially, a muscular *septum primum* grows inwards from the atrial roof (Anderson et al., 2003) associated with a mesenchymal cap at its leading edge and an additional mesenchyme at its base called the dorsal mesenchymal protrusion (Burns et al., 2016; Webb et al., 1998), and these fuse with mesenchyme at the AV complex (Deepe et al., 2020). Before its closure, the upper edge of the *septum primum* becomes fenestrated by apoptosis, forming a left-right communication termed the *ostium secundum* (Anderson et al., 2003; Moore and Persaud, 1998). An infolding of the atrial wall creates the *septum secundum* leaving an additional prominent inter-atrial communication termed the *foramen ovale* (Burns et al., 2016), completing the flap valve apparatus (Anderson et al., 2003) (**Figure 1A**).

**Figure 1.**
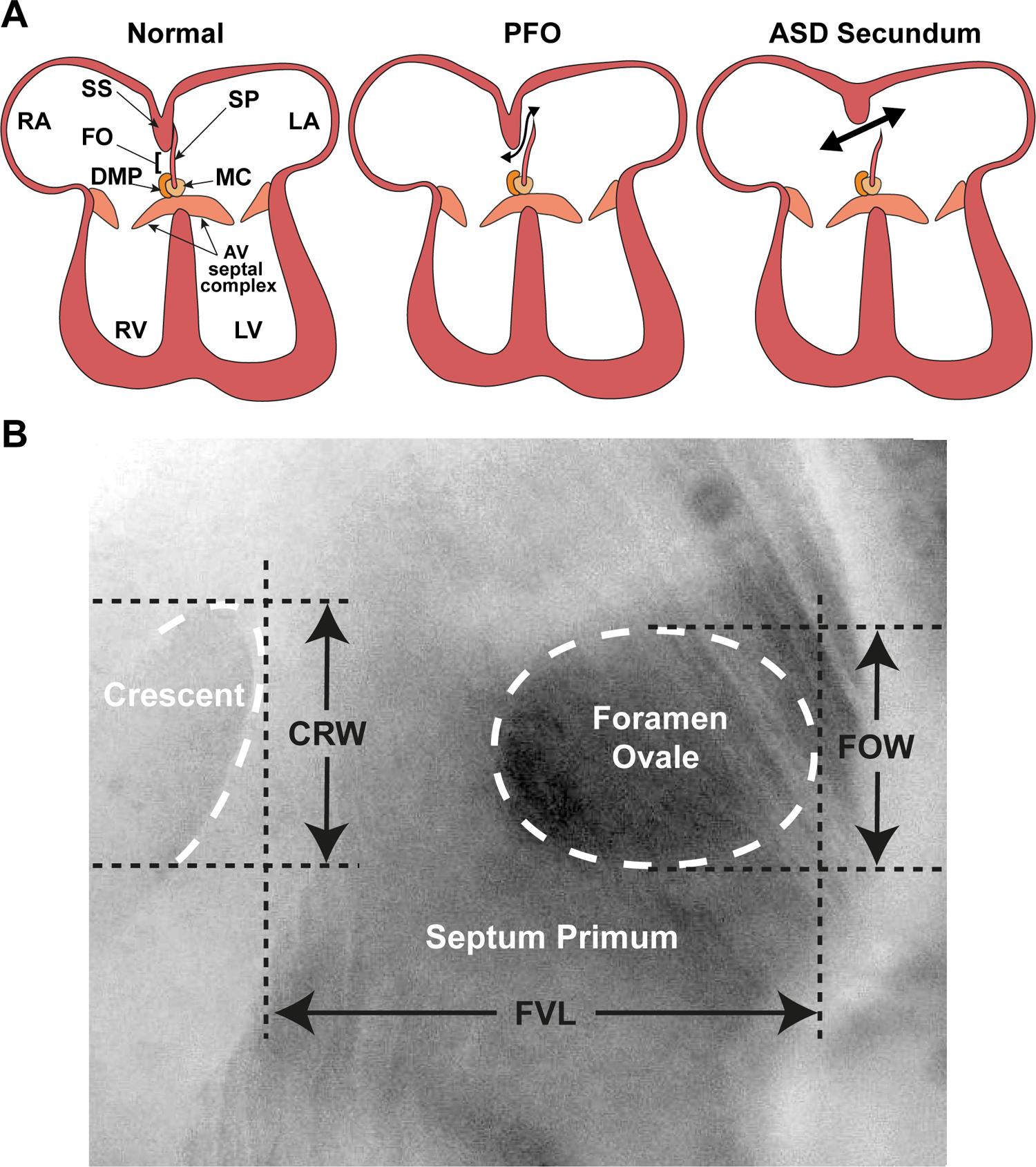
Atrial septal morphology in normal adult and congenital heart disease. **(A)** Schematic of atrioventricular septal complex and dysmorphology associated with patent *foramen ovale* (PFO) and more common secundum type of atrial septal defect (ASD). **(B)** Light micrograph of the interatrial septum of an adult mouse heart as seen after removal of the left atrial appendage. The annulus of the mitral valve is towards the lower left of this panel. Note that the *foramen ovale* is covered by the membranous atrial *septum primum*. Septal landmarks are shown along with quantitative atrial septal traits measured in F2 and F14 studies. The crescent corresponds to the upper edge (see **A**) of the atrial *septum primum*. Adapted from Kirk *et al*. 2006 (Kirk et al., 2006). AV: atrioventricular; CRW: crescent width; FO: *foramen ovale*; FOW: *foramen ovale* width; FVL: flap valve length; LA: left atrium; LV: left ventricle; RA: right atrium; RV: right ventricle; SP: *septum primum*; SS: *septum secundum*.

When the lungs become activated at birth, left atrial pressure increases and the flap valve normally closes permanently by fusion of the *septum primum* to the *septum secundum*. However, fusion is incomplete in about one-quarter of the human population, leading to the condition termed *patent foramen ovale* (PFO) (Hagen et al., 1984). For larger PFO, there is the strong probability of a hemodynamically significant inter-atrial communication associated with a higher risk of cryptogenic (unexplained) stroke, likely due to the passage of venous thrombi across the patent septum to the systemic circulation (Lechat et al., 1988; Webster et al., 1988). Larger PFOs are also associated with atrial septal aneurysm (ASA) (Hagen et al., 1984; Homma et al., 2003), as well as migraine with aura, clinical hypoxemia, and decompression illness in divers (Shnaider et al., 2004; Torti et al., 2004; Wilmshurst et al., 2000).

Atrial septal defect (ASD) is a less prevalent but more severe abnormality of the atrial septum (Feldt et al., 1971). There is likely a genetic link and anatomical continuum between the more common *secundum* form of ASD (ASDII) with PFO and ASA (Kirk et al., 2006; Kirk et al., 2007; Posch et al., 2010). ASDII arises from abnormalities of the *septum primum* and/or *secundum*, and presents as a frank and permanent interatrial corridor (Zipes, 2005), and, if untreated, the left-to-right blood shunt present postnatally can cause pulmonary hypertension and Eisenmenger syndrome, a life-threatening complication (Beghetti and Galie, 2009).

Analysis of quantitative trait loci (QTL) has emerged as an approach to understand the genetic complexity underpinning both quantitative and complex (non-Mendelian) binary traits (Ma et al., 2020; Shirai and Okada, 2021). Our previous study of inbred mouse strains revealed significant variation in atrial septal anatomy correlating with PFO and ASA (Biben et al., 2000), with genetic background as a major determinant. We established quantitative parameters of septal status including length of the *septum primum* (flap valve length; FVL), orthogonal width of the *foramen ovale* (FOW) and width of the open corridor in PFO (crescent width; CRW) (**Figure 1B**). We found that mean FVL was strongly negatively correlated with the prevalence of PFO across a variety of genetic backgrounds (Biben et al., 2000) and, collectively, short FVL, large FOW and large CRW were all strongly associated with PFO risk (Kirk et al., 2006). All traits were influenced by mutation of the cardiac homeodomain transcriptional master-regulator NKX2-5, pathogenic variants in which are strongly causative for ASDII in humans (Biben et al., 2000; Schott et al., 1998). We performed a QTL analysis with an F2 design using QSi5 and 129T2/SvEms strains representing mice with extremes of atrial septal phenotypes, identifying 7 significant (logarithm of odds (LOD) > 4.3) and 6 suggestive (LOD > 2.8) QTL affecting quantitative septal traits (Kirk et al., 2006), indicating a complex genetic basis for atrial septation defects in inbred mice. The F2 design used in our previous study has the power to detect the most significant QTL underlying complex traits, but confidence intervals are usually large, and each may reflect the effects of multiple loci and can contain hundreds of potential candidate genes (Darvasi et al., 1993). Therefore, increasing recombination has become the focus of fine mapping approaches including the use of advanced intercross lines (AIL), generated by intercrossing inbred strains with extreme phenotypes across 10 or more generations to increase chromosomal recombination (Darvasi and Soller, 1995). A similar rationale underlies the establishment of multi-parental recombinant inbred strains, which capture >90% of common genetic diversity among mouse species, used recently to study complex cardiovascular disease traits in adults (Salimova et al., 2019).

Here, we advance our understanding of genetic complexity underlying atrial septal variation in inbred QSi5 and 129T2/SvEms mouse strains. We used AIL to confirm and fine map significant QTL identified for FVL and FOW in the previous F2 study. We also sequenced the genomes of the AIL parental strains and integrated variant analysis with transcriptome data from dissected atrial septa at different developmental stages in parental strains. Our results provide the first high-resolution picture of genetic complexity underpinning atrial septal variation in the mouse model, allowing the identification of QTL with high impact, candidate genes and gene regulatory network perturbations that may have relevance to human PFO and ASD.

## Results

### Selection of AIL mice - atrial septal phenotypes

In our previous F2 study (Kirk et al., 2006), adult QSi5 and 129T2/SvEms parental mice and F2 mice were scored for PFO as a binary trait, and three quantitative anatomical parameters of the inter-atrial septum (FVL, FOW, and CRW) that were found to be associated with PFO (see Methods) (Biben et al., 2000; Kirk et al., 2006). The prevalence of PFO in parental strains was 4.5% and 80%, respectively. Beginning the pedigree from a single breeding pair, we randomly intercrossed F2 mice selected for extremes of phenotype for a further 12 generations to generate F14 (AIL) mice (see details in Methods). As in the original F2 study, the most relevant quantitative septal parameter in the F14 study was FVL (**Table 1**; **Figure 1-figure supplement 1**) - it showed the greatest difference in mean length up to a maximum of 2.5-fold between different inbred strains (Biben et al., 2000), and a difference of ∼2-fold and 4.8 standard deviations (SD) between parental strains for this study. FVL also showed the strongest (negative) correlation with PFO prevalence among a number of inbred strains (*r* = −0.97) (Biben et al., 2000), and in both F2 (*p* < 0.001) and F14 (*p* < 0.001) generations (Kirk et al., 2006).

**Table 1.** Phenotypic characteristics of parental strains and F2 mice extracted from Kirk (Kirk, Hyun et al. 2006) compared to F14 mice

|  | QSi5 | 129T2/SvEms | F2 | F14 |
| --- | --- | --- | --- | --- |
| N | 66 | 75 | 1437* | 933 |
| PFO (%) | 4.5 | 80 | 17 | 34 |
| FVL ± SD (mm) | 1.13 ± 0.11 | 0.60 ± 0.11 | 1.0 ± 0.19 | 1.01 ± 0.16 |
| FOW ± SD (mm) | 0.21 ± 0.06 | 0.24 ± 0.06 | 0.21 ± 0.07 | 0.24 ± 0.07 |
| CRW ± SD (mm) | 0.51 ± 0.13 | 0.44 ± 0.12 | 0.41 ± 0.12 | 0.54 ± 0.15 |
| Body Weight ± SD (g) | 29.4 ± 2.77 | 17.5 ± 2.1 | 26.6 ± 3.3 | 25.8 ± 2.9 |
| Heart Weight ± SD (g) | 0.21 ± 0.02 | 0.14 ± 0.02 | 0.21 ± 0.03 | 0.18 ± 0.03 |

\*Data were incomplete for some mice.

In our original study in which quantitative septal parameters were defined (Biben et al., 2000), we also determined the width of the patent corridor between the *septum primum* and *septum secundum* in cases of PFO, measured at the edge of the *ostium secundum* of the *septum primum* remnant (**Figure 1**), which forms a prominent crescent-shaped ridge. However, since this parameter was constrained to cases of PFO, for QTL analysis we considered only CRW (crescent width), defined as the length of the prominent crescent irrespective of the presence of PFO (Kirk et al., 2006). Whereas there was a strong statistical effect of PFO on CRW in the F2 study (*p* < 0.001) (Kirk et al., 2006), this was lost in the F14 cohort (*p* = 0.069) (**Supplementary File 1**). An additional observation was that mean CRW was positively associated with risk of PFO in the F2 cohort, whereas in the F14 study longer CRW was associated with lower PFO prevalence (negative association). For these reasons and because CRW remains an ill-defined anatomical parameter, we did not consider this trait in selecting mice with extremes of phenotype in either the F2 or F14 study. However, post-hoc analysis for CRW QTL in the F2 study revealed a significant QTL on MMU7 (LOD = 4.58) and a suggestive one on MMU3 (LOD = 3.49) (Kirk et al., 2006). We therefore performed a similar post-hoc analysis in this F14 study (see below).

FOW showed more subtle differences among individuals within the parental strains, substantially overlapping within one SD (**Table 1**), and the correlation with FVL was weak in the F2 cohort (*r* = − 0.087; *p* = 0.001) (Kirk et al., 2006). Nonetheless, variation in FOW is reflective of up to a 2-fold difference in *foramen ovale* area (Biben et al., 2000). Therefore, FOW was taken into account in selecting mice of extreme phenotypes for inclusion in the AIL study, and was indeed associated with both PFO (*p* < 0.001) (**Supplementary File 1**) and FVL (*r* = −0.284; *p* < 0.001) in F14 mice (**Supplementary File 2**).

In the AIL study, we sought to confirm and fine map significant QTL found in the F2 study with LOD scores above the threshold for significance of 4.3. This included three QTL for FVL and three for FOW. We also included a suggestive QTL (2.8 < LOD < 4.3) for FOW located on MMU9 (LOD = 3.43), since its peak covered the T-box transcription factor gene *Tbx20*, mutations in which are known to cause familial septal defects and severe PFO (Kirk et al., 2007).

### Selection of AIL mice - heart weight and body weight phenotypes

It was evident that the quantitative septal parameters under study might be influenced by heart size and mass (**Table 1**) (Kirk et al., 2006), and indeed both FVL and FOW were significantly correlated with HW in both F2 and F14 cohorts, albeit that the effects were small (**Supplementary File 2**) (Kirk et al., 2006) and HW had no influence the likelihood of PFO. In the F2 study, therefore, we did not normalize septal data for HW so as not to mask important QTL (Kirk et al., 2006). However, the possibility that FVL and FOW QTL could be explained by variation in HW or BW has not been formally excluded. Thus, prior to analysis of AIL mice, we performed a retrospective linkage analysis for HW and BW on F2 data. The HW of F2 mice was initially adjusted for factors with significant effects (age, sex, and BW) and we used the same LOD score criteria as in the F2 study (4.3 for significant and 2.8 for suggestive linkage) (Lander and Kruglyak, 1995). We discovered a suggestive QTL (LOD = 3.4) for normalized HW on MMU7, and another on the same chromosome that fell just short of suggestive (**Figure 2A**). The suggestive HW QTL overlapped with previously determined QTL for BW on MMU7, but did not represent a BW QTL in our study. We also found a significant QTL for HW normalized for age, sex and BW on MMU11 (LOD = 8.5) (**Figure 2B**). This QTL overlapped one for BW normalized for age and sex (LOD = 14.2) (data not shown). Thus, we included normalized HW as a parameter in selection of F14 mice with extremes of phenotype for further QTL analysis (see Methods), and selected markers for fine mapping of the significant normalized HW QTL on MMU11.

**Figure 2.**
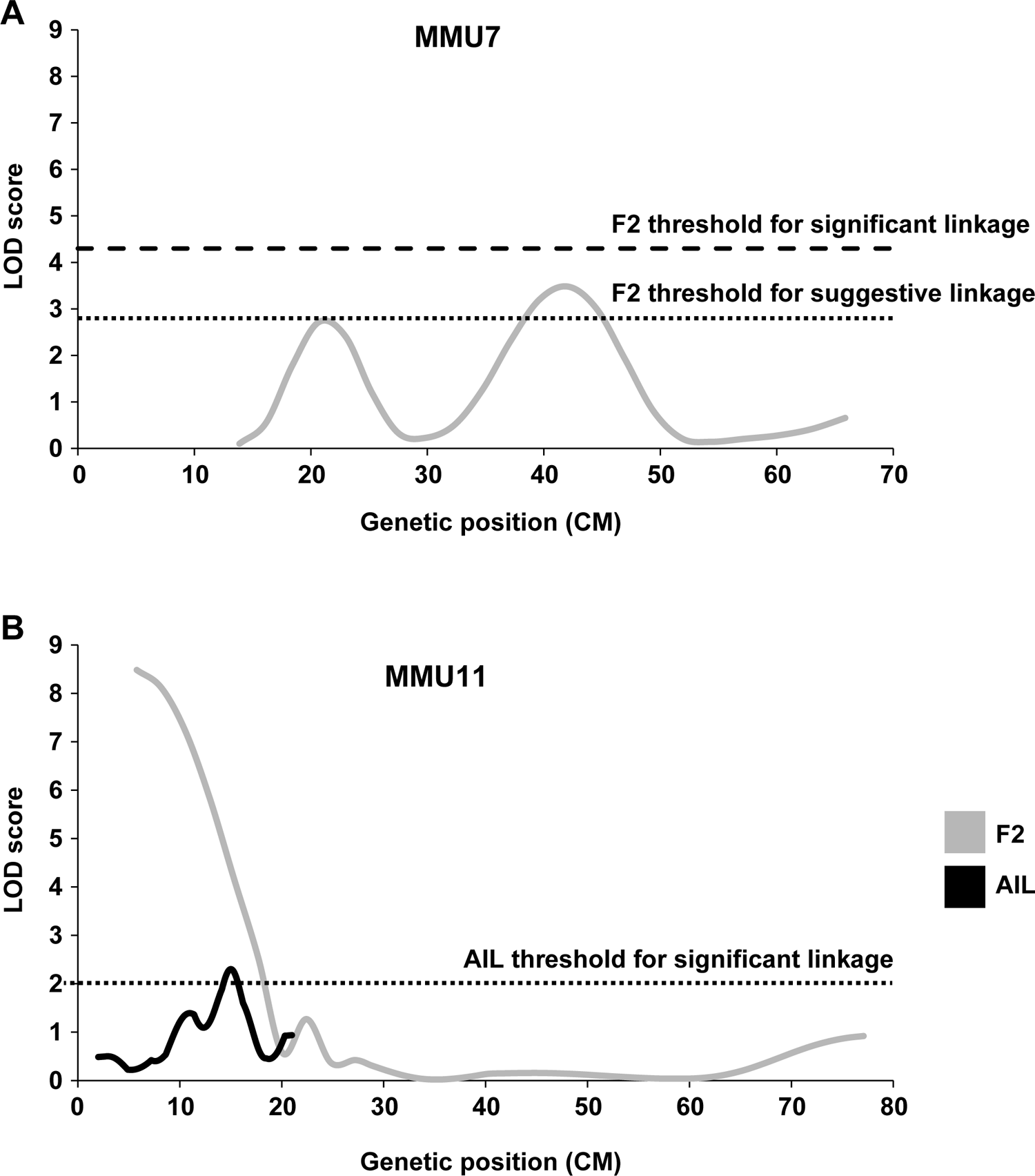
Linkage analysis for HW. Suggestive and significant QTL identified by the F2 study. **(A)** With the latter being fine mapped by the AIL study. **(B)** Y-axes represent LOD scores for HW adjusted for age, sex, and BW.

### Linkage results for atrial septal morphology

We set a LOD score of 2 as a cut-off for significant linkage based on the density of chosen markers (∼2 cM) and the size of the genomic regions covered (Lander and Botstein, 1989). From the septal morphology QTL identified in the F2 study (six significant and one suggestive), at least six QTL were confirmed and significantly narrowed using AIL data. Five F2 QTL resolved into multiple peaks and several new QTL were also discovered. Furthermore, the overlap between QTL for different traits was increased. **Supplementary File 3** describes QTL for FVL, FOW, and CRW identified by the AIL study and **Figure 3** shows the examples from each relevant chromosome comparing F2 and AIL mapping results. Overall, 37 QTL were significant in the AIL study. As in the F2 study, the direction of QTL varied, and were classified here as “normal” if they contribute to the negative septal features documented for the 129T2/SvEms strain (shorter FVL; longer FOW; shorter CRW) and “cryptic” if protective (**Supplementary File 3**).

**Figure 3.**
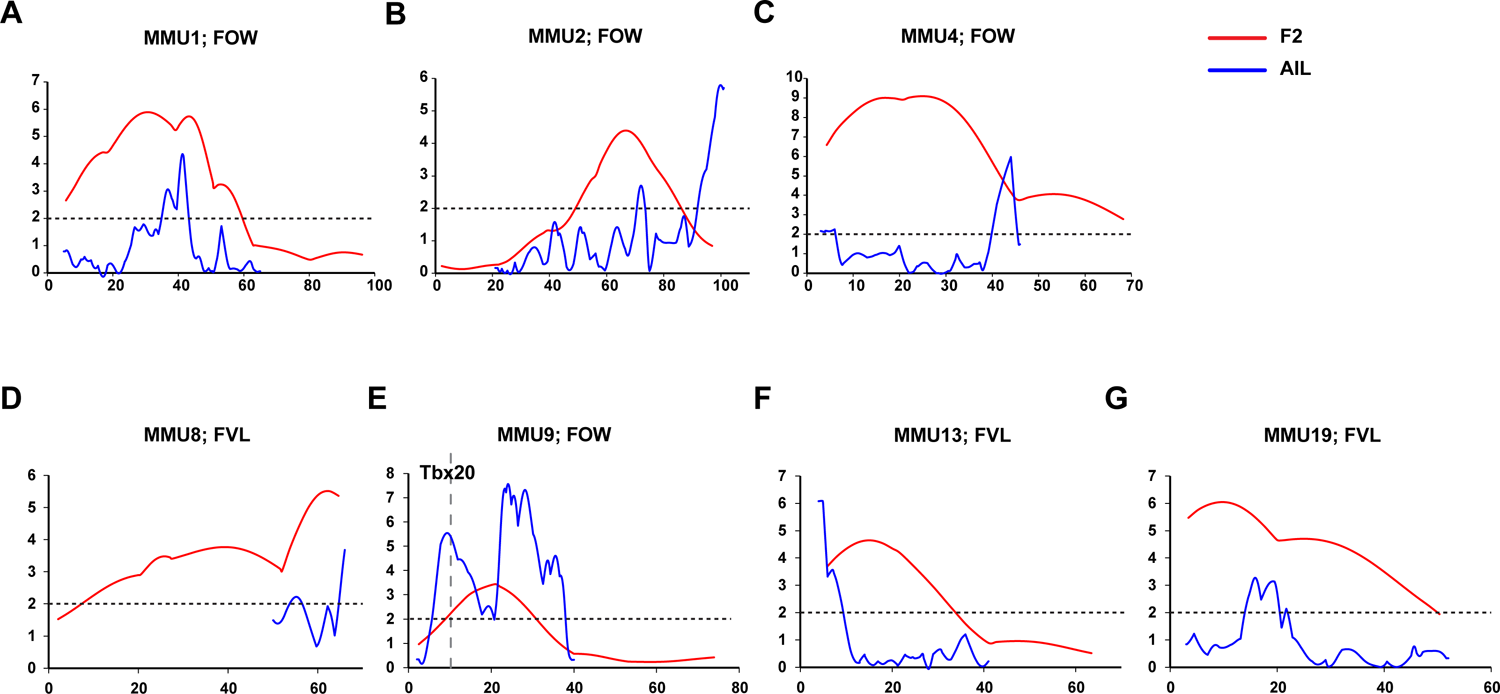
Comparison of linkage results from F2 (gray line) and AIL (black line) populations. Y-axes represent LOD scores and x-axes represent genetic map positions. A 1 LOD drop-off for each QTL is shown on the x-axes representing the confidence interval of the QTL. SNP markers within AIL QTL are also shown.

### Linkage analysis of PFO as a binary trait

Although in general the linkage analysis of continuously distributed traits is more sensitive and informative than analysis of binary traits, the analysis of PFO as a binary trait was still of importance in our study. Therefore, we also performed direct linkage analysis for the presence or absence of PFO in the F14 mice as a binary trait using a logistic regression model that we previously developed (Moradi Marjaneh et al., 2012). This analysis confirmed most of the FVL and FOW QTL. The AIL results for each chromosome for quantitative (FVL, FOW, and CRW) and binary (PFO) atrial septal traits are compared in **Figure 3**. Results from each chromosome are summarized as follows:

### MMU1

We previously identified a significant QTL for FOW with a peak at 30.8 cM extending 26.1 cM (inclusive of one LOD on each side of the peak = 1-LOD drop-off) on MMU1 (**Figure 3A**). The linkage analysis of AIL data narrowed this region to 7.5 cM, including two adjacent peaks with LOD scores of 3.1 and 4.4 overlapping in the 1-LOD drop-off. To assess whether the two peaks were distinct, we re-ran the linkage analysis including the marker closest to the distal peak as a fixed term (**Figure 3-figure supplement 1A**). While this led to a significant reduction in LOD score of the distal peak, the proximal peak did not change and was still significant, indicating that the peaks represent two independent QTL. Unlike the F2 study, the AIL study also showed strong evidence of linkage across this region for FVL (including two FVL QTL with LOD scores of 6.0 and 2.3 respectively) (**Figure 4A**). Analysis of the AIL also revealed an additional QTL for FVL on MMU1 located at 9 cM, with peak LOD of 2.2, which is clearly separate from the original QTL (**Figure 4A**). The QTL map from analysis of PFO as a binary trait followed the pattern produced by FOW and FVL data (**Figure 4A**). In particular, its peak at 41.6 cM (LOD = 4) strongly supported highly significant FOW and FVL QTL in that vicinity. The 1-LOD drop-off of this peak covered approximately 2.4 cM, a very substantial refinement on the F2 study.

**Figure 4.**
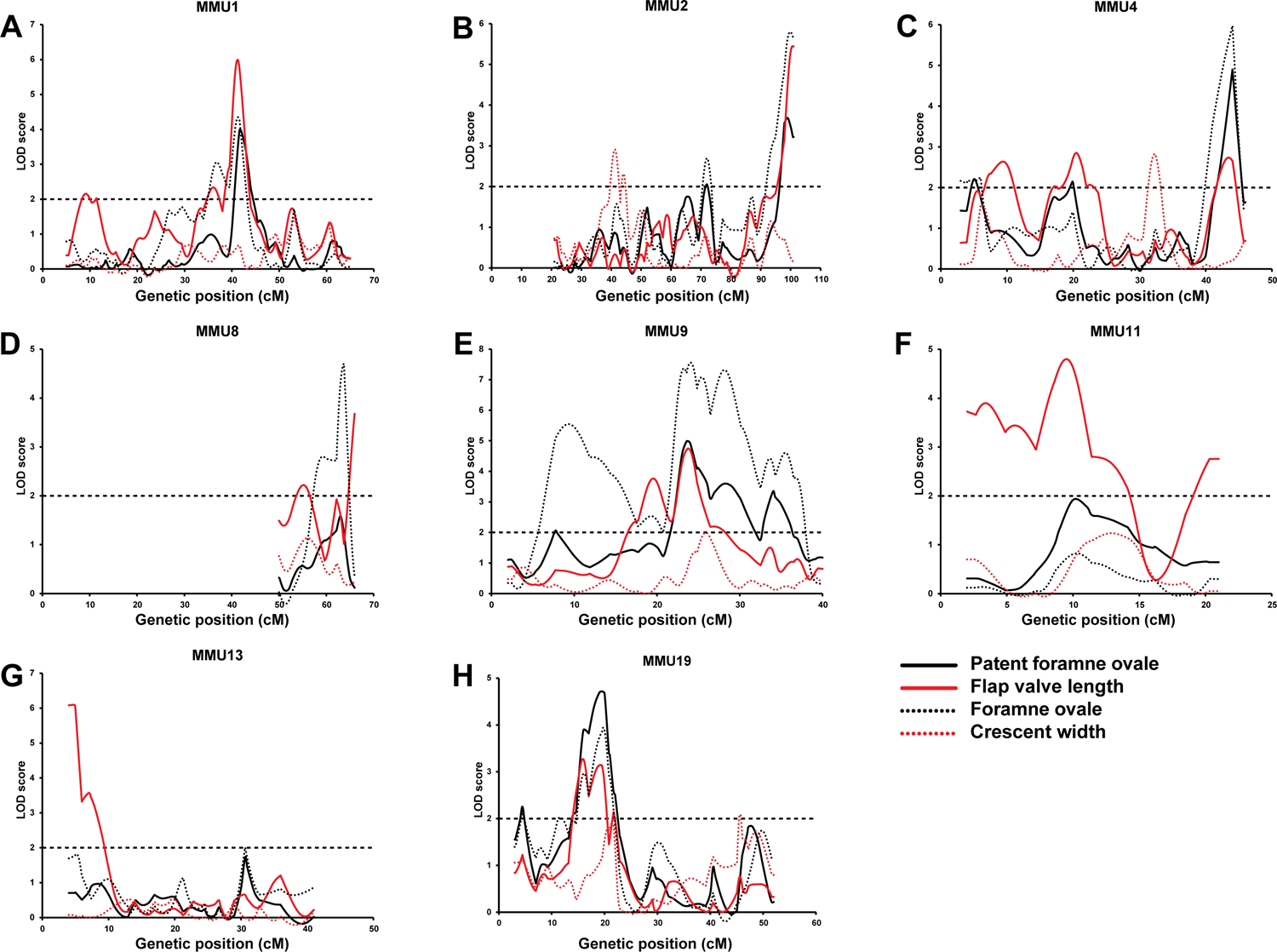
Comparison of AIL results for PFO, FVL, FOW, and CRW. Y-axes represent LOD scores and x-axes represent genetic map positions. Vertical blue lines correspond to the positions of the AIL markers.

### MMU2

The broad F2 QTL peak for FOW (peak at 67.5 cM; **Figure 3B**) was narrowed to a sharp peak at 72 cM refining the QTL genomic region from 18.2 cM (F2) to 3.6 cM (AIL) (**Figure 3B** and **Figure 4B**). We also observed a new highly significant QTL at the distal end of this chromosome (∼100 cM) affecting both FOW and FVL (LOD scores 5.79 and 5.44, respectively) (**Figure 3B** and **Figure 4B**). The binary analysis of PFO supported QTL for both FOW and FVL. While the F2 study was not able to detect QTL for CRW on this chromosome, linkage analysis of the AIL data revealed two possibly distinct QTL close to each other at 41.25 cM and 44 cM (**Figure 4B** and **Figure 3-figure supplement 1B**) without pleiotropic effects on the other quantitative traits.

### MMU4

The very broad F2 FOW QTL spanning 23.8 cM of the chromosome (peaking at 25.4 cM; **Figure 3C**) resolved in the AIL as two FOW QTL (6 cM, LOD = 2.2; 44 cM, LOD = 6.0), leaving the mid portion including the 1-LOD drop-off of the original peak unlinked (**Figure 3C**). Given the sparsity of markers across this region in the F2 study, this outcome likely represents a resolution of the broad F2 QTL region into two widely spaced QTL. The AIL data also revealed three QTL for FVL, of which the distal one was located at the same position as the distal FOW QTL (**Figure 4C**). The analysis of PFO as a binary trait generated a similar pattern to the FVL and FOW results. A new QTL for CRW was detected at 32.25 cM which, as for the CRW QTL on MMU2, did not affect the other traits.

### MMU8

The region at the end of this chromosome was found to be linked to FVL in the F2 study (**Figure 3D**). It peaked at 62.5 cM and covered 12.2 cM of the chromosome. On analysis of the AIL data, this QTL resolved into two QTL at 52.25 cM and 66 cM (**Figure 3D**). A highly significant QTL was also detected for FOW at 63.7 cM (**Figure 4D**).

### MMU9

The F2 study revealed a suggestive QTL for FOW peaking at 20.7 cM (LOD = 3.4) and extending 17.3 cM on MMU9 (**Figure 3E**). The AIL resolved this into at least four separate QTL with peaks at 9.25, 19.25, 24, and 28.25 cM (**Figure 3E**). *Tbx20*, a cardiac transcription factor gene, is located at 10.25 cM, within the 1-LOD drop-off of the first QTL and very close to its peak at 9.25 cM. We also detected a new QTL for FOW, peaking at 35.5 cM with maximum LOD score of 4.6. Analysis of the AIL also identified two peaks for FVL at 19.5 cM and 23.75 cM with significant LOD scores (**Figure 4E**), both overlapping with significant peaks for FOW. The analysis of PFO as a binary trait resulted in a strikingly similar pattern to the FOW, significantly supporting all 4 of the FOW QTL, including the QTL coinciding with the *Tbx20* gene.

### MMU11

The F2 study did not show linkage of any region of MMU11 to the atrial septal traits. However, this chromosome was included in the AIL study to fine map the HW QTL detected by the analysis of the F2 data (see below). Using the chosen markers for the HW QTL we also discovered at least two new QTL underlying FVL, with peak LOD score of 4.8 at 9.5 cM (**Figure 4F**).

### MMU13

The AIL narrowed down the broad genomic region of the F2 QTL for FVL (**Figure 3F**) from 19.4 cM to 8.7 cM (**Figure 4G**). It is notable that the AIL data resulted in a shifting of the QTL peak from 15.3 cM (F2) toward the telomere of the chromosome including two close peaks at 4.9 and 7 cM (AIL) which were determined to be significantly distinct using the fixing method (see Methods) (**Figure 3-figure supplement 1C**).

### MMU19

The broad F2 QTL for FVL at 10.2 cM resolved into three separate QTL with the highest LOD score at 16 cM (**Figure 3G**). The fixing method showed these peaks represented three separate QTL (**Figure 3-figure supplement 1D**). We also observed linkage between this region and FOW, with a highest LOD score of 3.9 at 19.75 cM (**Figure 4H**). Binary analysis of PFO showed a similar pattern to that seen for FVL and FOW, strongly supporting the QTL located between 13 cM and 23 cM.

### Linkage analysis for heart weight

As noted above, our retrospective analysis of F2 data revealed a significant QTL for normalized HW at the proximal end of MMU11, with highest LOD score of 8.5, and a suggestive QTL on MMU7 peaking at 42.1cM (**Figure 2**). This is the first report of QTL affecting HW on MMU11, although numerous HW QTL on other chromosomes have been detected previously using different cohorts and study designs (Reed et al., 2008; Rocha et al., 2004; Sugiyama et al., 2002). An extra 9 markers covering the MMU11 HW QTL region were selected for this AIL study (see Methods). We also performed linkage analysis for normalized HW on other chromosomal regions for which markers had been previously chosen for analysis of atrial septal morphology. MMU2, MMU4, MMU9, MMU11, and MMU13 all showed significant evidence of linkage to HW (**Figure 2-figure supplement 1**); of these, only MMU2 has previously been linked with HW, albeit on different genetic backgrounds (Rocha et al., 2004). On MMU11, we observed evidence of linkage distally with a LOD score of 2.3 peaking at 15 cM, thus confirming the original F2 QTL (**Figure 2B**).

Comparison of linkage results for HW and atrial septal morphology, however, showed that only the HW QTL on MMU2 and MMU9 had a potential overlap within its 1-LOD drop-off with QTL for atrial septal parameters. Thus, the chromosomal locations of QTL for HW were largely different from those defining atrial septal parameters.

### Contribution of QTL to phenotypic differences

The percent attributable phenotypic variance for each QTL is shown in **Supplementary File 3**. As in the F2 study (Kirk et al., 2006), many AIL QTL were of relatively large effect - 18/37 individually contributed ∼23-70% of the difference between parental means. Interestingly, the majority of these (15/18) were for FOW, with 6 being normal and 8 being cryptic QTL. Normal and cryptic QTL are anticipated to interact additively or in more complex ways in parental strains and individual mice, which may show phenotypes more extreme than parental strains (Kirk et al., 2006). The effect sizes of FVL QTL were somewhat smaller, individually 0.3-9% of the difference between parental means, with the majority (12/15) being normal QTL. Whereas it is difficult to precisely quantify the effects of individual or combinations of QTL, our data suggest that many of the QTL detected are of large or moderate effect size and therefore collectively contribute significantly to quantitative trait variation between parent strains.

### Whole genome sequencing of parent strains and SNP density analysis

The whole genomes of the AIL parental strains were sequenced and mapped to the C57Bl/6J mouse reference genome (NCBI38/mm10). Genomic analysis of the various inbred mouse inbred strains used in research reveal a mosaic pattern of haplotypes attributable to the genome of founder lines. Accordingly, pairwise comparisons of inbred strain genomes show long blocks of either low or high SNP density representing shared and divergent ancestry, respectively (Frazer et al., 2007; Wade et al., 2002). Regions with a high rate of polymorphism between parental strains (spanning nearly one-third of the genome but harboring more than 95% of the genetic variation) can significantly reduce the regions of interest underlying QTL (Wade et al., 2002), albeit that regions with low SNP density may contain rare variants that have arisen since the establishment of parent strains (Bloom et al., 2019).

High-quality homozygote variants between the AIL parental strains identified by whole genome sequencing were used to compute variant density across the genome (**Figure 5-figure supplement 1**; see Methods). To assess the distribution of variant densities, we binned the genome into non-overlapping intervals and counted the number of variants in each bin (**Figure 5-figure supplement 2**). The analysis was repeated using different bin sizes ranging from 50 kb to 1.1Mb. The distribution of variant densities across the genome showed a bimodal pattern, confirming two categories of genomic regions with low and high variant density regions. Binning at 600kb intervals showed the clearest distinction between the two peaks of distribution and the boundary between the two peaks, representing a cut-off of 1000 variants per 600 kb interval, was chosen to classify genomic segments as low or high variant density regions. Intersecting with high variant density regions narrowed the total size of the QTL regions from 264.1Mb to 109.5Mb (41.5%) (**Figure 5**). It is noteworthy that among the high variant density regions called there were blocks of very high variant density, presumably also derivative of strain ancestry.

**Figure 5.**
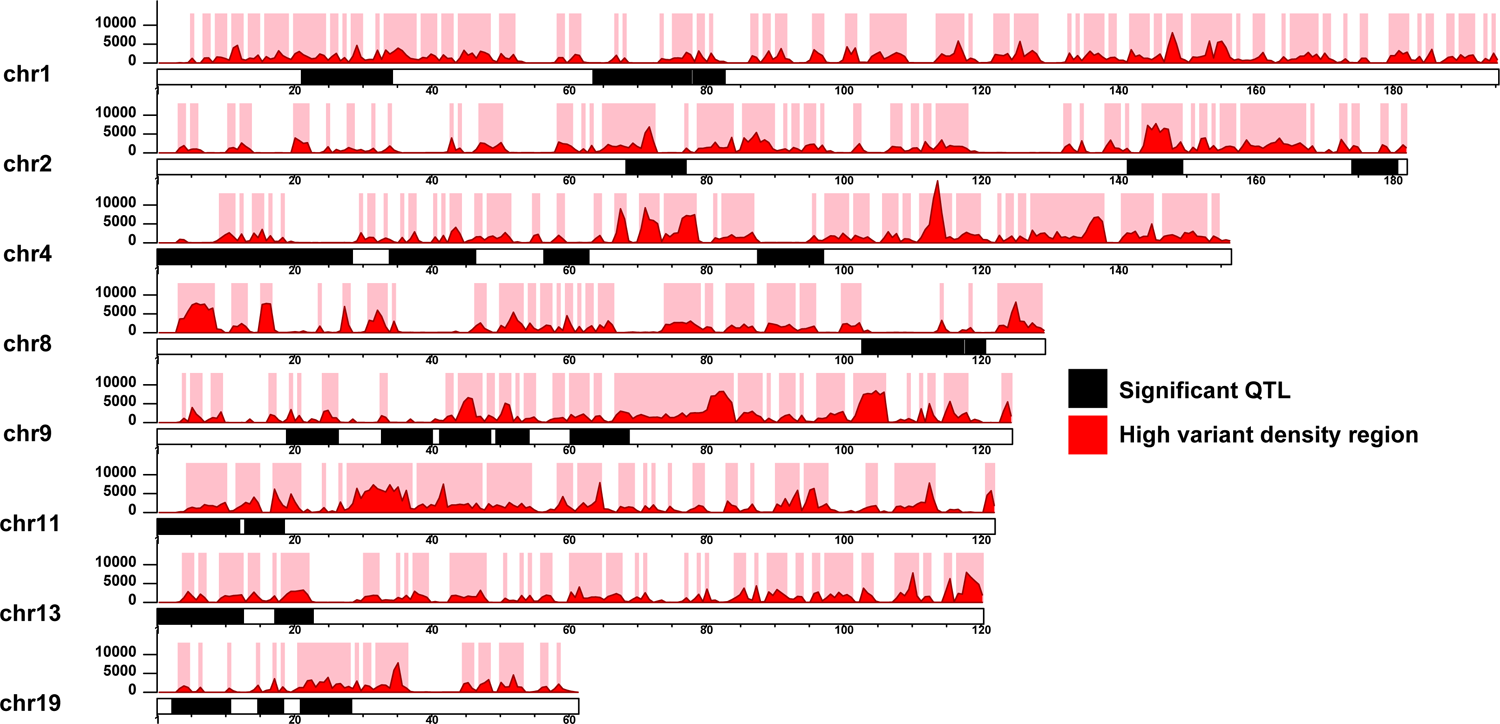
Narrowing down the QTL using high variant density regions. The QTL regions for each chromosome are illustrated as black blocks. The red histograms and y-axes represent the variant density defined as the number of 129T vs QSi5 variants per 600Kb genomics intervals. High variant density regions (density ³ 1000) are represented as pink blocks. The x-axis for each chromosome represents the genomic position (Mb).

### Filtering for high impact variants and candidate genes

There were a total of 6,769,034 high quality variants between the AIL parental strains identified by whole genome sequencing (using C57Bl/6J as reference). We first filtered for protein-coding variants in genes expressed in the developing atrial septum (see below) and which lay under QTL regions (1-LOD drop-off support interval), then analysed through the Ensembl Variant Effect Predictor (VEP) tool (McLaren et al., 2016) for pathogenicity impact. We filtered variants for high impact (stop-gain, stop-loss, start-loss, splicing defect, frameshift), missense variants predicted deleterious by SIFT (Ng and Henikoff, 2003) and non-high impact (e.g. non-frameshift) deletions. This resulted in 45 variants spread over 28 genes for 129T2/SvEms and 47 variants across 35 genes for QSi5 (**Supplementary File 4**; **Figure 6A,B**), noting that two genes had unique variants in each parent strain (total number of genes carrying predicted deleterious variants = 61). Virtually all of these were in high SNP density regions and many genes (26%) contained multiple variants (**Figure 6A,B**). Only synonymous variants were found in *Tbx20*. To create a high confidence QTL gene list, we annotated genes according to prior association with cardiovascular phenotypes (Mouse Genome Informatics; MGI) or link to human CHD from a curated list of high-confidence and emerging CHD genes (Alankarage et al., 2019). A total of 27/61 QTL genes (44%) containing high impact or non-synonymous protein-coding variants overlapped with one or both lists distributed between both parent strains (**Figure 6C,D**). We highlight the following examples of predicted deleterious and high confidence genes:

**Figure 6.**
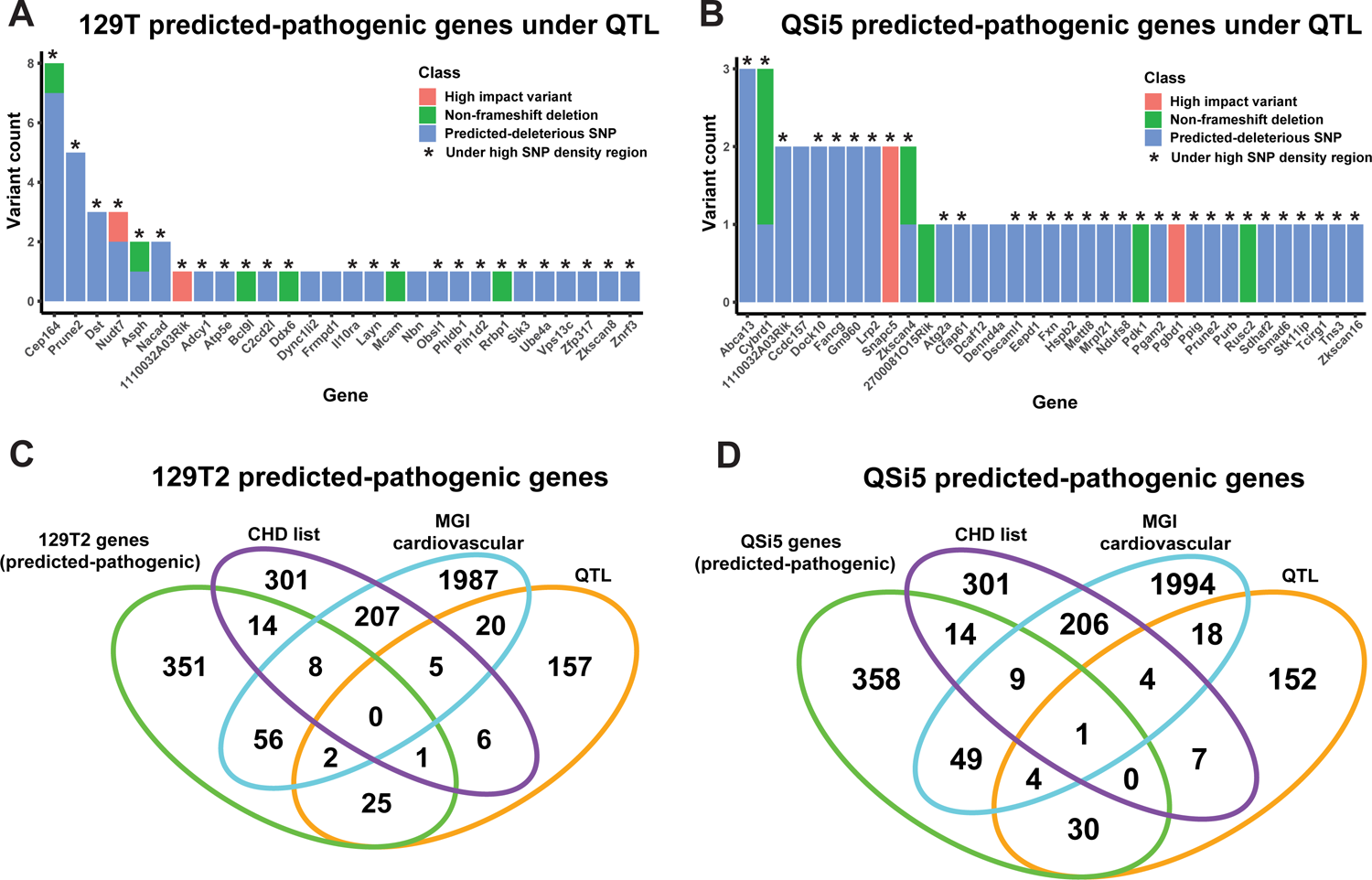
Identification of predicted-pathogenic variants. **(A,B)** Number of variants per gene according to classification (High-impact, non-frameshift deletion or predicted-deleterious missense variant) for **(A)** 129T2 and **(B)** QSi5. **(C, D)** Overlap of genes containing predicted-pathogenic variants with 1) genes in a high confidence/emerging CHD list, 2) genes with a heart development cardiovascular function from MGI and 3) all genes under QTL intervals that contain protein-modifying variants. Overlaps are shown for **(A)** predicted pathogenic variants in **(C)** 129T2 and **(D)** QSi5.

#### Cep164

In the 129T2/SvEms strain under a cryptic FOW QTL on MMU9, *Cep164* contained seven predicted-deleterious missense coding variants and one in-frame 3 base pair (bp) deletion (**Figure 6A**). *Cep164* encodes a docking protein associated with mother centrioles and is essential for formation of primary and multi-cilia, therefore affecting cellular processes such as fluid sensing and inter-cellular signalling (Schmidt et al., 2012). Defects in ciliogenesis are well known to cause CHD (Audain et al., 2021; Li et al., 2015). Three *Cep164* variants overlap regions predicted to encode coiled coil domains in the centre of the protein and one variant may overlap a predicted DNA/RNA-binding domain (PredictProtein (Bernhofer et al., 2021)). *Cep164* knockout mice are embryonic lethal showing multiple severe defects including heart developmental arrest at the looping stage (Siller et al., 2017).

#### Dst

In the 129T2/SvEms strain under a cryptic FVL QTL on MMU1, *Dst1* carried three missense mutations predicted to be deleterious. *Dst1* encodes diverse isoforms of Dystonin, members of the plakin family of cytoskeletal docking and tumour suppressor proteins implicated in cell growth, adhesion, cytoskeleton organisation, intracellular transport and migration. They may function in part through an ability to antagonise activation of the Hippo pathway effector YAP (Jain et al., 2019). Mutations in humans and mice cause fragile skin (epidermolysis bullosa), severe neurological conditions, as well as skeletal muscle instability (Horie et al., 2017; Kunzli et al., 2016). Isoform BPAG1b is expressed in cardiac and skeletal muscles colocalising with Z-bands, sarcolemma and cardiac intercalated discs, and interacts with α-actinin2. Knockout strains show diminished heart size and atrophy before weaning and altered hypertrophy/wall stress markers (Horie et al., 2017). *Dst* is classified as an emerging CHD gene.

#### Sik3

In the 129T2SvEms strain under a cryptic FOW QTL on MMU9, *Sik3* carries a non-synonymous SNP predicted to affect SIK3 function. *Sik3* encodes a serine/threonine kinase of the AMP Kinase (AMPK) family whose activity is inhibited by cyclic AMP (cAMP) via phosphorylation at multiple sites by cAMP kinase (PKA). SIK3 functions in a variety of biological processes (Wein et al., 2018) including through phosphorylation and inhibition of cAMP-regulated transcriptional coactivators (CRTCs) and class IIa histone deacetylases (HDACs), the latter functioning as inhibitors of MEF2 transcription factors that play key roles in heart development (Darling and Cohen, 2021). *Sik3* overlaps with the MGI list with a homozygous targeted EUCOMM mutation showing increased heart weight.

#### Lrp2

in the QSi5 strain under a CRW QTL on MMU2, *Lrp2*, encoding a low density lipoprotein receptor-related protein, carried two missense variants predicted to be deleterious. *Lrp2* is expressed in the SHF progenitor pool and developing outflow tract (OFT), and is known to act in multiple morphogenetic pathways. Variants in human *LRP2* have been associated with hypoplastic left heart (Theis et al., 2020), whereas mutant mice show premature differentiation and depletion of anterior SHF progenitors in the fetus, leading to a shortened OFT, *truncus arteriosus*, and ventricular wall and septal defects (Christ et al., 2020).

#### Smad6

Under a cryptic FOW QTL on MMU9 in the QSi5 strain, *Smad6*, encoding an inhibitory member of the *mothers against decapentaplegic* transcription factor (SMAD) family, carried the non-synonymous SNP NM_008542 c.845C>G (p.Arg282Pro), predicted to be deleterious. We chose this variant for further analysis. SMAD6 is an inhibitor of BMP signalling acting at multiple levels, including inhibition of the binding of receptor-activated (R) SMADs 1, 5 and 8 (the effector transcription factors of the BMP pathway) to BMP receptors, where they are normally activated by phosphorylation; inhibition of binding of R-SMADs to their co-SMAD (SMAD4); and direct transcriptional repression of targets genes of R-SMADs by recruiting transcriptional repressors such as CtBP, HDAC-1/3 and Hoxc8/9 (Bai and Cao, 2002; Bai et al., 2000; Lin et al., 2003) (**Figure 7A**). BMP signalling has been implicated in cardiac specification and development of cardiac chambers, valves, septa and outflow tract (Wang et al., 2011), and SMAD6 mutations in humans are associated with syndromic and non-syndromic congenital malformations, including cardiac outflow tract defects, as well as patent *foramen ovale* and atrial and ventricular septal defects (Calpena et al., 2020; Galvin et al., 2000; Tan et al., 2012). The R282P variant lies within the regulatory linker region joining the MH1 DNA-binding domain and MH2 transactivation/protein:protein interaction domain (**Figure 7B**), and interrupts the conserved PPXY motif known to bind the E3 ubiquitin ligase SMURF1, predicted to poly-ubiquitinate SMAD6 triggering proteosome-mediated degradation (Sangadala et al., 2007). SMURF1 ubiquitinates other targets including SMAD1, and can also act to exclude SMAD1 (and possibly SMAD6) from the nucleus (Sapkota et al., 2007). The R281P variant also lies close or immediately adjacent to serine residues that are modifiable by phosphorylation (Bian et al., 2014; Christensen et al., 2010; Mertins et al., 2016). In SMAD1, SMURF1 binding and subsequent ubiquitination is dependent upon MAPK-mediated phosphorylation of linker serine/threonine residues, which also prime phosphorylation by kinase GSK-3β, facilitating poly-ubiquitination (Sapkota et al., 2007). Thus, R282P SMAD6 variant potentially disrupts the complex regulatory network regulating SMAD6 stability and activity.

**Figure 7.**
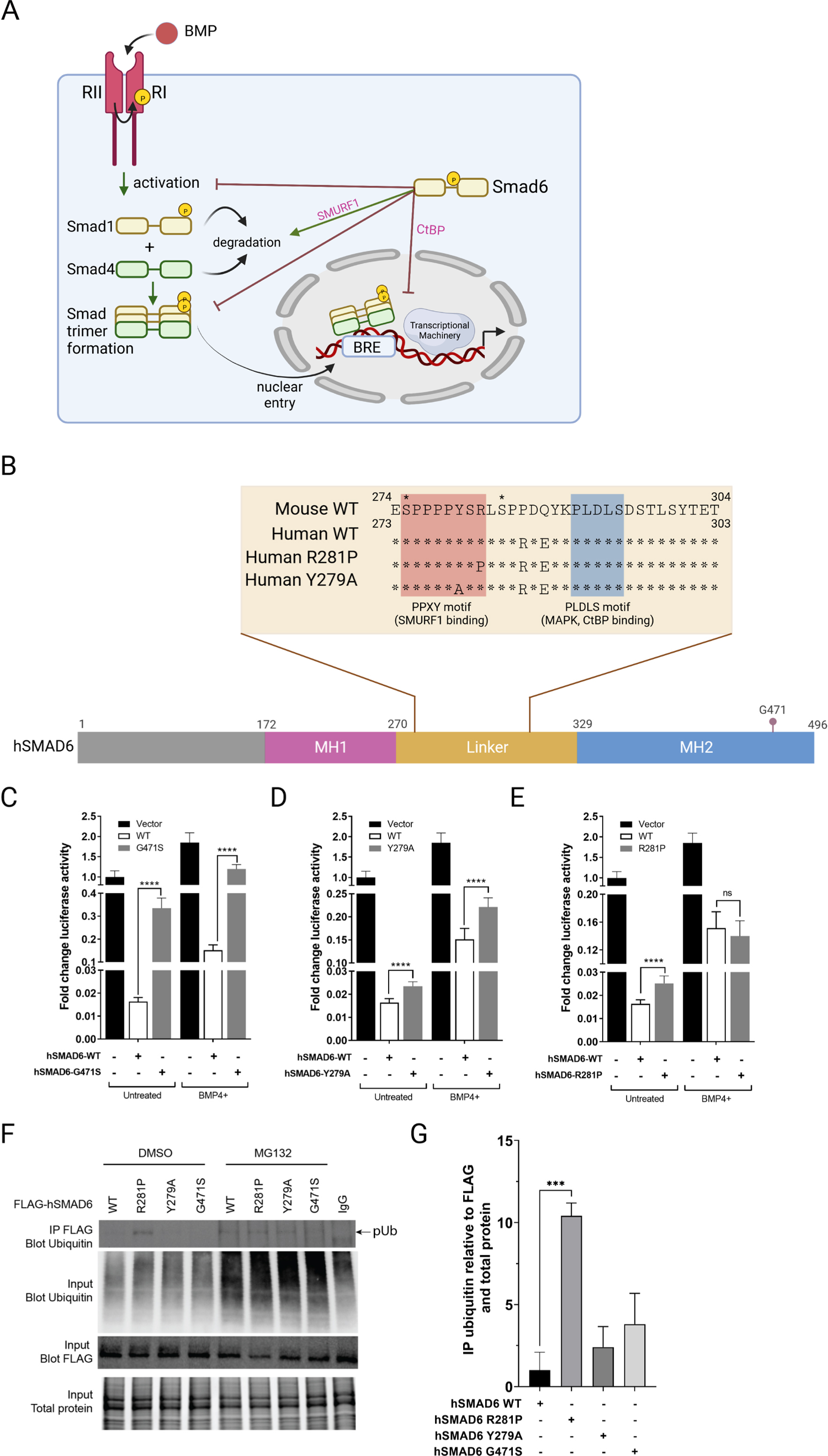
Functional analysis of SMAD6 variant R282P. **(A)** Schematic of BMP signalling pathway highlighting roles of SMAD6. **(B)** Schematic of hSMAD6 showing its three main functional domains - the conserved DNA-binding MH1 domain (172-270aa), regulatory linker region (271-328aa), and the C-terminal transactivation/cofactor interaction MH2 domain (329-496aa). The position of G471 within the MH2 domain is indicated – mutation of this residue (G471S) disrupts the interaction between SMAD6 and SMAD1. Insert shows amino acid sequence of the linker region with the arrowhead indicating R282 (considered wild type) found in C56Bl/6 and 129T2/SvEms strains. The PPXY (SMURF1 binding) and PLDLS (MAPK, CtBP binding) domains are indicated, along with sequences of hSMAD R281P (equivalent to R282P as in the mouse QiS5 strain) and Y279A variants. Known serine phosphorylation sites are marked * above the sequence. **(C-E)**. Functional analysis of hSMAD6 variants. Activity of hSMAD6-G471S **(C)**, hSMAD6-Y279A **(D)**, and hSMAD6-R281P **(E)**, were assessed relative to wild type (WT) hSMAD6 on a BMP-signalling responsive (BRE)-luc promoter in untreated and BMP4-treated cells. n=6-9; differences between WT and variant were assessed by unpaired t-test; data presented as mean ± SD. **(F,G)** Ubiquitination of WT hSMAD6 and variants assessed in MG132 or DMSO (control)-treated HEK293T cells. WT FLAG-ShMAD6 and variants were immunoprecipitated with anti-FLAG antibody with lysates subject to western blotting with anti-ubiquitin antibody **(F)**. Total ubiquitin, FLAG-hSMAD6 and protein indicated. **(G)** Quantification of FLAG-hSMAD6 ubiquitination. n=3; difference between WT and variants assessed by unpaired t-test; data presented as mean ± SD.

We first analysed the activity of WT and mutant forms of SMAD6 in HEK293T cells after transfection of human (h) cDNAs and a luciferase reporter construct reading out SMAD-dependent BMP-signalling (**Figure 7C-E**). WT hSMAD6 repressed BMP signalling by ∼98% at baseline and ∼92% after stimulation of cells with BMP4. Repression was diminished significantly by the hSMAD6 mutation G471S, which abolishes the interaction between SMAD6 and SMAD1 (**Figure 7C**). Then, to gauge the impact of loss of interaction with SMURF1, we replaced tyrosine 279 of the hSMAD6 PPXY domain, predicted to be essential for SMURF1 binding, with alanine (Sangadala et al., 2007). Relative to WT, the Y279A variant slightly reduced SMAD6 repression at baseline, and the change was greater after BMP4 stimulation (**Figure 7D**). The R281P hSMAD6 variant (equivalent to R282P in mouse), also showed a slight reduction in repression similar to Y278A at baseline, however, the reduction was eliminated after BMP4 stimulation (**Figure 7E**), suggesting an alternative mechanism to simply loss of SMURF1 binding. To take a different approach, we tested the poly-ubiquitination status of the above hSMAD6 variants using co-immunoprecipitation (**Figure 7F**). After transfection of FLAG-tagged hSMAD6 vectors into HEK293T cells and treatment with proteosome inhibitor MG132 or DMSO control, lysates were precipitated using a FLAG antibody and ubiquitination of hSMAD6 detected by western blotting. In the presence of MG132, poly-ubiquitinated hSMAD6 WT, R281P and Y279A was readily detected (**Figure 7F**). Ubiquitination of G471S was lower, perhaps because ubiquitination depends on interaction with an R-SMAD. In DMSO, poly-ubiquitination of WT, Y279A and G471S hSMAD6 were barely detected; however poly-ubiquitination of R281P occurred at levels ∼10-fold higher than WT, comparable to that seen in the presence of MG132 (**Figure 7F,G**). Our findings suggest a defect in the SMAD6 R281P-SMURF1 binding/ubiquitination pathway leading to constitutive poly-ubiquitination.

### Transcriptome analysis of the developing atrial septum

To explore changes in the gene regulatory networks affecting atrial septal traits, we profiled the transcriptome of developing septa dissected from AIL parental strains at embryonic stages (E) 12.5, E14.5, and E16.5 (*n*=6/sample; see Methods). RNA was extracted and pooled to create two replicate pools per mouse strain per time point (three septa/pool; total 12 samples) and cDNA libraries were generated and sequenced on an Illumina platform.

Principal component analysis (PCA) performed on the top 500 variable protein-coding genes showed that sample replicates clustered closely together, whereas samples from the different embryonic time points segregated along the first principal component (PC1) axis and those from the different inbred strains segregated along the PC2 axis (**Figure 8A**). PCA performed independently on annotated expressed non-coding genes also showed separation of embryonic time points and strains (**Figure. 8B**). We calculated differentially expressed protein coding genes (DEGs; protein coding) for each time-point separately (using the DESeq2 R package (Love et al., 2014)) using significance thresholds of p_adj_ < 0.05 and log2 fold-change > 0.5. E12.5 had the largest number of differentially expressed genes (990), followed by E16.5 (881) and E14.5 (762) (**Figure 8C,D**; **Supplementary File 5**) and at all time-points there were more genes down-regulated in 129T2/SvEms compared to QSi5 (**Figure 8C**). The majority of DEGs were unique to a time-point; however, 453 genes were down-regulated in at least 2 time points (**Figure 8D**).

**Figure 8.**
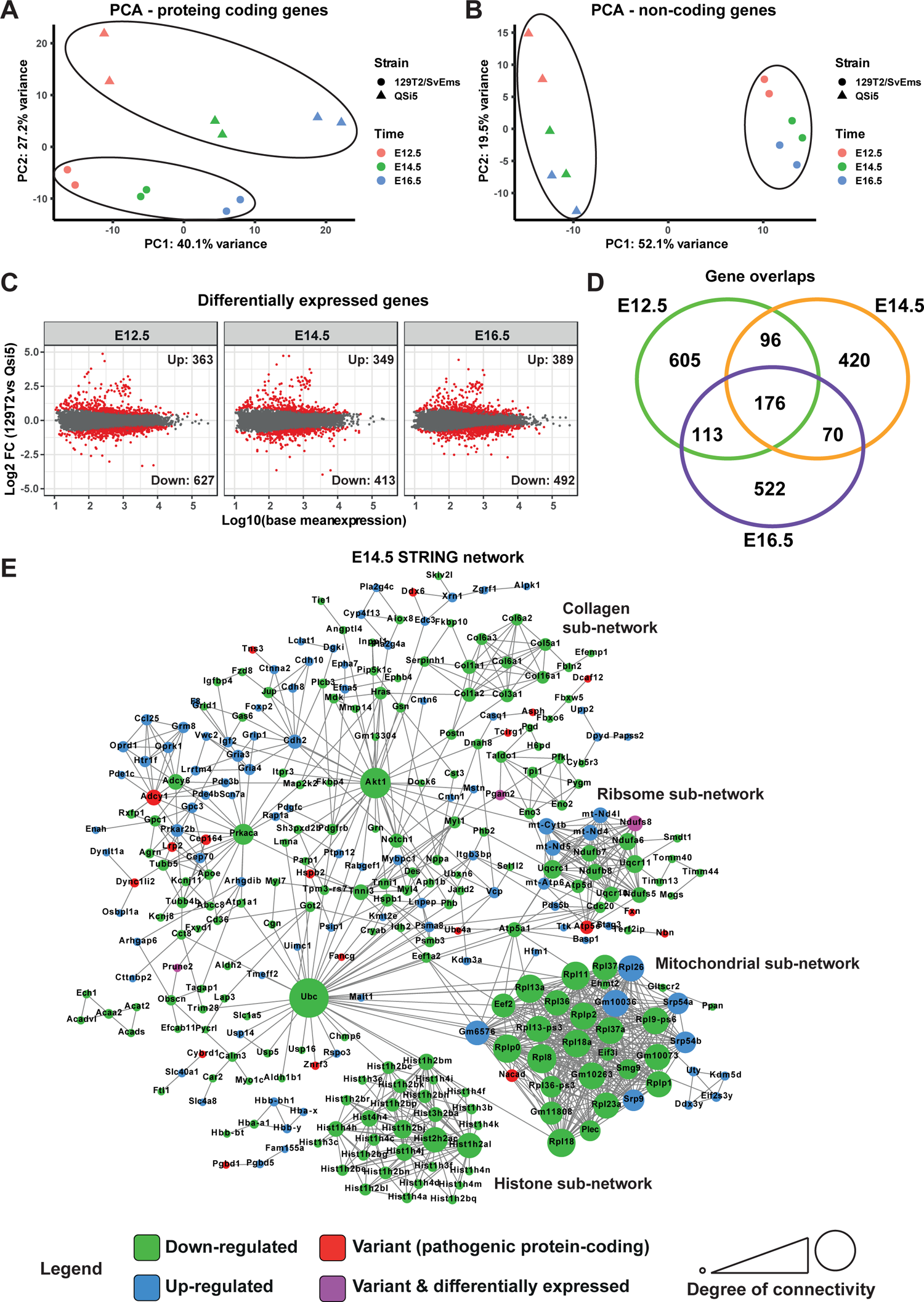
Transcriptome analysis of atrial septum. (**A, B**) Principal component analysis (PCA) plot of gene expression profiles from atrial septum RNA-seq libraries including two replicates per time point (E12.5, E14.5, and E16.5) per mouse strain (QSi5 and 129T2/SvEms) for **(A)** protein coding genes and (**B**) non-coding genes. (**C**) MDA plots of differentially expressed genes for each time point. (**D**) Venn diagram with the number of differentially expressed genes between QSi5 and 129T2/SvEms at different time points. (**E**) STRING network of E14.5 differentially expressed genes and genes with predicted-pathogenic variants from QSi5 or 129T2/SvEms. Genes are coloured according to whether they are upregulated or downregulated in 129T2/SvEms, contain a predicted-pathogenic variant, or are both differentially expressed and containing a variant.

We investigated the biological significance of DEGs at each time point separately using protein-protein association connections obtained from the STRING database (Szklarczyk et al., 2017), filtering for high confidence connections. Overall network significance was calculated by comparing network metrics to those from STRING networks of randomly selected genes across 100,000 permutations. To discover potential drivers of network perturbations, we also included in the STRING analysis protein coding genes which were located under QTL (1 LOD drop-off) and in which variants predicted to be deleterious had been identified. Using this stringent approach, the network at E14.5 was significant for both the number of edges (observed = 980; expected = 492.8±63.2; *p* < 1×10^-5^) and average clustering coefficient (observed = 0.39; expected = 0.28±0.03; p = 0.0001) (**Figure 8E**). The E14.5 network was driven predominantly by genes downregulated in 129T2/SvEms compared to QSi5 septa, i.e. the strain with highest prevalence of PFO and most disadvantageous septal traits (shortest FVL, largest FOW). Highly connected sub-networks contained genes involved in biosynthetic and signalling pathways and macromolecular machines, including nucleosomes, ribosomes, mitochondria and extracellular matrix (ECM) (**Figure 8E**). This signature was supported by significant over-representation of gene ontology (GO) biological process terms among DEGs for *nucleosome assembly, chromatin assembly, translation, ATP metabolic process and extracellular matrix organization* (**Supplementary File 6**; Fisher’s exact test; false discovery rate < 0.05). Whereas networks for E12.5 and E16.5 alone, and those for DEGs common across all time points, were not significant overall, both E12.5 and E16.5 networks contained ribosomal and ECM protein sub-networks (**Figure 8-figure supplement 1 and Figure 8-figure supplement 2**).

Within the E14.5 network, Ubiquitin C (*Ubc*) was a prominent down-regulated hub gene connecting directly to histones (specifically H2B) and ribosomal genes as well as others dispersed across the network, most of which were also down-regulated (**Figure 8E**). In addition to its well-characterised role in protein homeostasis, *Ubc* is also involved in transcription and genome integrity (Mark and Rape, 2021; Mattiroli and Penengo, 2021; Qu et al., 2021). As such it might be expected to be functionally connected to a large number of genes even in random networks; however, the number of connections between *Ubc* and other genes in the septal network was far greater than expected by chance (Fisher’s exact test, p_adj_ = 8.8×10^-5^). Furthermore, analysis of network topology in Cytoscape showed that *Ubc* had the highest measure of *stress centrality* and *betweenness centrality*, key indicators of network hubs (Shannon et al., 2003). Among high confidence QTL genes carrying predicted deleterious variants were *Znrf3* and *Ube4a*, encoding ubiquitin conjugation enzymes (E3 and E4 ligases, respectively), providing a potential link between pathogenic variants and transcriptome output.

Other genes with high centrality scores were *Akt1* and *Prkaca*, encoding AKT Serine/Threonine kinase/Protein Kinase B (AKT/PKB) and Protein Kinase A (PKA), respectively, which were down-regulated in 129T2/SvEms compared to QSi5 septa, and had dispersed down-regulated connections across the network. These connections were over-represented in their respective known and predicted phosphorylation targets (sourced from PhosphoSitePlus (Hornbeck et al., 2012) and PhosphoPICK (Patrick et al., 2015)) - 12/32 for ATK, 10/19 for PKA (*p* < 1×10^-4^ for both kinases; Fisher’s exact test) (**Figure 8E**). We further analysed known and predicted kinase targets among all genes in the network, which resulted in an additional 35 phosphorylation connections for PKA and 9 for AKT. Genes identified as lying under QTL and carrying predicted deleterious variants were also identified as AKT and/or PKA targets, including *Cep164* (see above) and *Atf2*, encoding a Zn^2+^ multi-functional transcription factor and chromatin modifier involved in cAMP-dependent transcriptional response (**Figure 9**).

**Figure 9.**
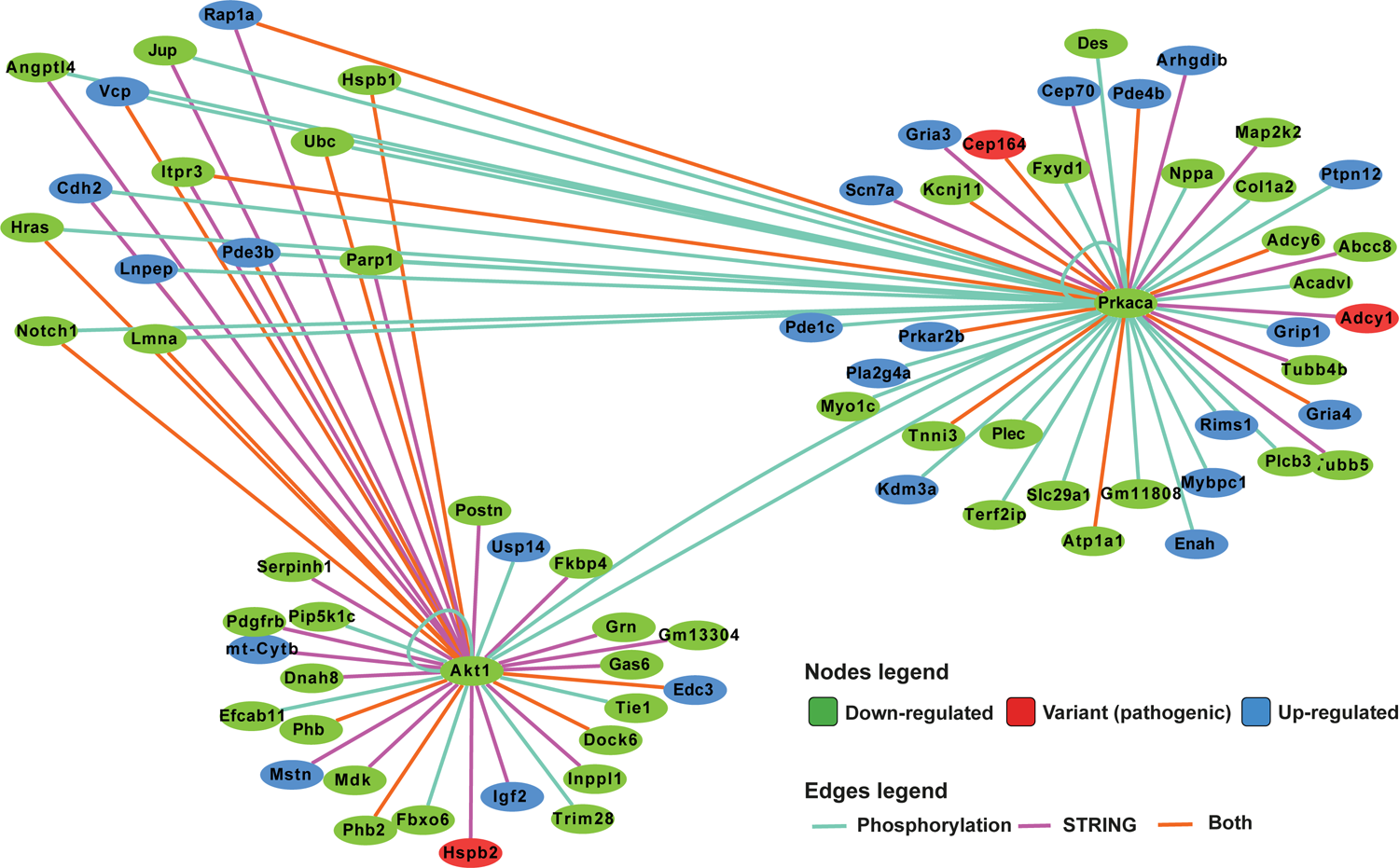
Network analysis of PKA and AKT1 phosphorylation targets. Integrated network analysis of PKA and AKT1 STRING connections and phosphorylation substrate targets from PhosphoSitePlus and PhosphoPICK predictions.

### Differentially expressed QTL genes are enriched for genomic variants

Both protein coding and *cis*-regulatory variants are likely causally linked to network perturbations underlying quantitative traits, as highlighted by GWAS on human disease (Zhang et al., 2014). To explore the relationship between QTL variants and DEGs further, we assessed enrichment of variants in and around DEGs underlying QTL relative to their genomic features (enhancers, promoters, 5’UTRs, exons, introns and 3’UTRs), comparing to variant distribution in non-DEGs. Enhancers were defined as regions showing diagnostic enhancer histone (H) marks (H3K4me^+^ or H3K27ac^+^; H3K4me3^-^) within 50kb upstream of the transcriptional start site of relevant genes. We selected DEGs as protein coding genes with an adjusted differential expression *p*-value (*p*_adj_) of <0.05 and absolute log2 fold-change of >0.5 in at least one atrial septal time-point. Of 2,168 such genes, 214 genes were under QTL (1 LOD drop-off). For non-DEGs, we selected 2,168 protein coding genes which showed the highest adjusted *p*-values after merging data from all three time-points, of which 188 genes were under QTL. After permutation testing, we observed a significant enrichment of 129T2/SvEms vs QSi5 variants relative to expected in most genomic features (enhancers, promoters, 5’UTRs, exons, and introns) for DEGs underlying QTL, whereas non-DEGs showed a significant depletion of variants (**Figure 10**). DEGs and non-DEGs showed similar variant count distributions across features, albeit that non-DEGs had an increased likelihood of bearing zero variants (**Figure 10-figure supplement 1**), overall indicating that DEGs were not substantially skewed towards genes with high variant density. Furthermore, variants were not focused on any specific gene class (**Supplementary File 7).** These data suggest a strong mechanistic link between variant architecture and differential expression of genes under QTL.

**Figure 10.**
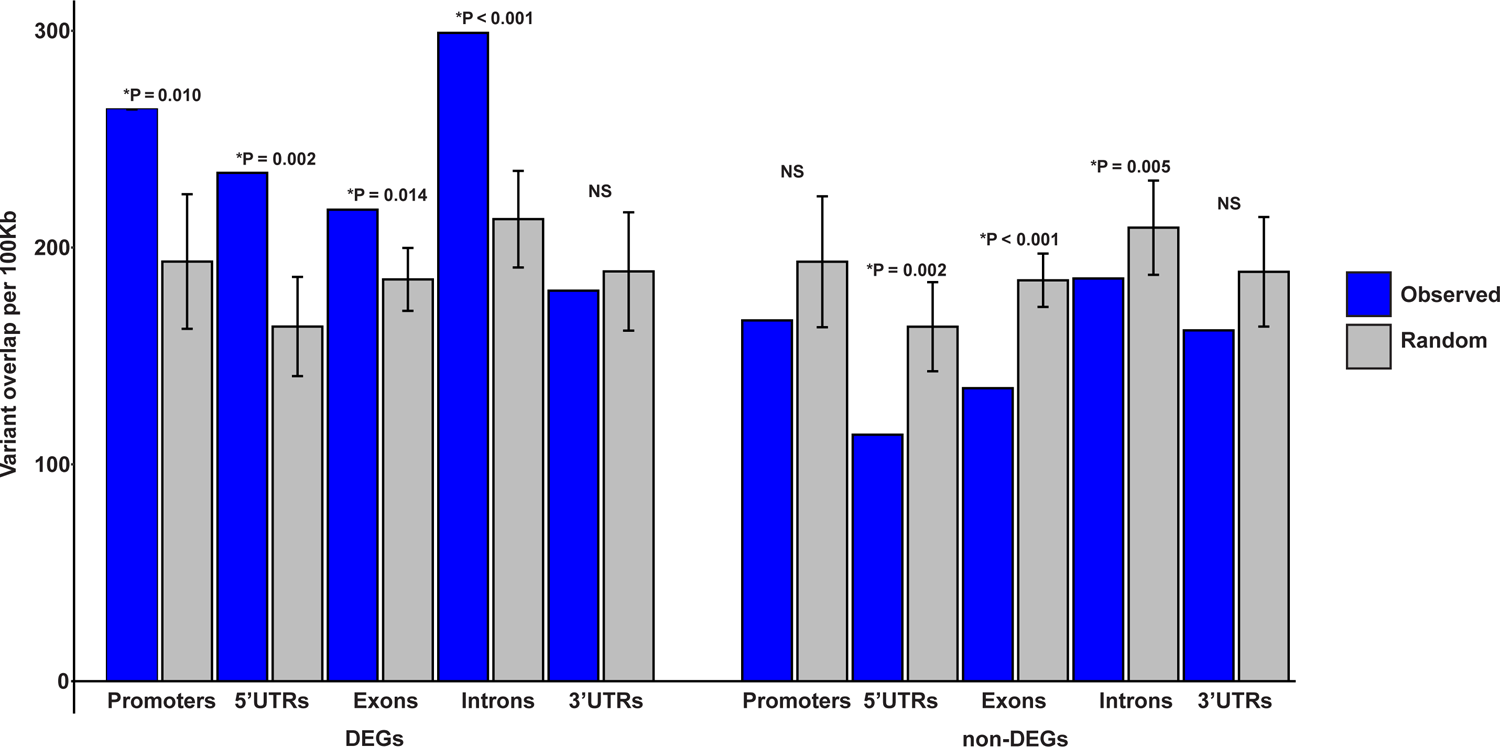
Enrichment of 129T2/SvEms vs QSi5 variants in genomic features of DEGs under QTL. The number of variants directly overlapping a feature is shown in blue, and gray bars represent the expected values based on the mean overlap from 1000 randomly generated interval sets. Error bars show the 95% confidence intervals of the mean. The significance of the enrichment is expressed as P-values, calculated by dividing the number of random samples showing equal or greater overlap than the observed by the total number of permutations (\**p* < 0.05).

## Discussion

QTL analysis has emerged as an approach to understand the genetic complexity underpinning both quantitative and complex (non-Mendelian) binary traits. PFO and ASD are examples of complex binary traits of medical significance. One model for complex binary traits assumes an underlying continuous but unobservable variable (termed liability) with a threshold above which an individual expresses a phenotype (Falconer, 1965). Quantitative parameters act as proxies for the assumed liabilities and significantly increase the power of QTL detection. Comparing F2 and AIL designs with identical parameters including sample size and marker density, the predicted confidence intervals of AIL QTL is *t*/2 times smaller than those of F2 QTL, where *t* is the number of AIL generations (Darvasi and Soller, 1995). This indicates that AIL is a powerful method for precise localization of QTL and also separation of linked QTL identified by an F2 design.

We previously mapped QTL underlying quantitative parameters of the atrial septum using an F2 intercross design and here applied the AIL approach for confirmation and fine mapping. The cascade breeding program generating the F14 AIL resource from 48 breeding pairs per generation came close to the practical optimum (100 animals produced per generation) (Darvasi and Soller, 1995). From seven F2 QTL included in our AIL study, at least six were confirmed and substantially narrowed, whereas five F2 QTL resolved into multiple peaks and additional QTL were discovered. Overall, 37 significant atrial septal QTL were discovered.

Among the three quantitative parameters studied here, FVL and FOW showed a similar pattern of linkage on most of the chromosomes. This is a new finding that was not evident in the F2 study. Furthermore, independent analysis of PFO as a binary trait strongly supported results for FVL and/or FOW on most chromosomes. As noted, of the traits analysed, FVL has a larger variation between parental strains (**Table 1**) (Kirk et al., 2006) and shows a stronger negative correlation to PFO prevalence among several inbred and mutant strains (Biben et al., 2000), as well as PFO risk in both F2 and F14 mice. Collectively, these data suggest that many QTL affect formation of the primary and secondary atrial septa in common, and that FVL is a robust indicator for atrial septal morphology and risk of PFO.

CRW did not show a significant correlation to PFO in the AIL data and was therefore not found to be a satisfactory surrogate for the size of the open corridor in cases of PFO (Biben et al., 2000). Nonetheless, we found 5 significant QTL for CRW, only one of which overlapped with QTL for the other traits, suggesting that CRW is determined by largely different genetic elements to those governing FVL and FOW. As CRW is measured along the lower boundary of the apoptotic domain that generates the *ostium secundum* in septal development, it may relate to tissue remodelling subsequent to the apoptotic process.

As septal parameters may be influenced by heart size, we performed a linkage analysis for normalized HW across MMU11, where a significant QTL was discovered retrospectively in the F2 data, and indeed across all other chromosomal regions for which markers were selected. We confirmed and refined the position of the HW QTL on MMU11, and detected HW QTL on MMU2, MMU4, MMU9, and MMU13. Of these, only MMU2 has previously been linked with HW (Rocha et al., 2004). Intertrait correlation and linkage results showed that HW influences quantitative septal parameters in both F2 and F14 studies, but only in a minor way (**Supplementary File 2**). In addition, HW and atrial septal morphology showed limited overlap. We propose that HW is mostly determined by ventricular chamber growth as an independent parameter from the growth of “primary” myocardial components of the early heart tube, which have a low proliferative index (Moorman and Christoffels, 2003), and additional mesenchymal and cushion components that contribute to formation of the inter-atrial septum.

Overall, we conclude that septal morphology and risk of PFO has a complex genetic basis in inbred strains of mice. Furthermore, of 37 QTL for atrial septal morphology, many had effect sizes of ∼5-70% of the difference between parent strains, demonstrating the power of the AIL approach to discern alleles of major impact. Given that the analysis was restricted to the previously found QTL and the selected markers covered only a limited part of the genome, it is likely that the genetic complexity underpinning septal defects in the two inbred strains under study is even greater than revealed here. It is noteworthy also that the two inbred lines in this study were previously selected for independent traits. For example, the QSi5 strain, used here because of its low PFO risk (Kirk et al., 2006), was bred for numerous traits related to high fecundity (Holt et al., 2004) and whereas this has facilitated the generation of >1000 mice required for the AIL study, the genetic diversity that contributes to variation in atrial septal morphology may also be limited by prior selection. High genetic complexity underpinning variation in atrial septal morphology may be expected, given the diverse lineage origins, tissue contributions, and morphogenetic network processes contributing to septal structure (Anderson et al., 2003; De Bono et al., 2018; Deepe et al., 2020; Rana et al., 2014; Steimle et al., 2018). Extrapolating to the diverse outbred human population, we might conclude that potentially many hundreds of gene variants contribute significantly to atrial septal dysmorphology.

To create a list of protein coding genes that could potentially contribute to differences in atrial septal morphology in inbred strains of mice, we sequenced the whole genomes of parent strains then filtered for genes which lie under QTL (1 LOD drop-off), were expressed in the developing inter-atrial septal septum, and carried variants predicted to be pathogenic. The total list comprised 61 genes carrying 92 variants, resulting in an average ∼2.5 candidate protein coding variants per QTL, or ∼1.6 candidate deleterious protein coding genes per QTL. By far most of the predicted pathogenic variant genes lay within regions of high SNP density and 26% of them carried multiple variants. To create a higher confidence list, we further filtered for genes that overlapped the MGI list of mouse genes associated with cardiovascular phenotypes, and a curated known and emerging human CHD gene list (Alankarage et al., 2019). Our high confidence list contained genes involved in diverse developmental and cellular functions, including those encoding kinases, transcription factor, metabolic enzymes, membrane receptor/adhesion molecules, RNA helicase, cilial building block, and ubiquitin pathway elements. The role of most of these genes in septal development has not been elucidated. Our follow-up study of SMAD6 variant R282P demonstrated a hyper-ubiquitinated state for this critical regulator of BMP signalling. The full mechanism and *in vivo* significance remain to be determined, noting that hyper-ubiquitination, in addition to affecting SMAD6 stability *in vivo*, could inhibit additional regulatory processes or protein:protein interactions occurring throughout the linker region (Sapkota et al., 2007; Xu et al., 2016).

Extending genome analysis, we determined the transcriptome of atrial septal regions of both strains at three developmental timepoints. By far the most significant DEGs were at E14.5 and related to macromolecular structures including nucleosomes, ribosomes, mitochondria and ECM, with virtually all associated genes downregulated in the 129T2/SvEms strain, which shows less robust septal parameters. STRING networks identified three potential hub drivers of this state - genes for Ubiquitin c, and kinases AKT and PKA, with many of the genes encoding targets of these kinase being up- or downregulated also. We propose that the net impact of variants in parent strains affects development (e.g. SMAD, AKT, PKA), proteostasis (UBC), and biosynthetic and metabolic pathways impacting differential septal growth and morphogenesis independently of heart weight.

Our analysis of variant architecture under QTL (**Figure 10**) showed that variants (129T2/SvEms vs QSi5) were overrepresented in differentially expressed atrial septal genes located under QTL across multiple gene features including enhancers, promoters, 5′-UTRs, and introns. Non-DEGs showed underrepresentation of variants in most of these features. These data suggest a mechanistic link between the presence of non-coding (including cis-regulatory) as well as coding variants in the differential expression and function of septum genes underlying QTL, between parent strains. These genes are candidate drivers of network perturbations revealed by STRING analysis. A substantial proportion of both predicted-pathogenic genes under QTL (26%; **Figure 6**) and regulatory regions in DEGs under QTL (29%; **Figure 10-figure supplement 1**) bear multiple variants. Therefore, we predict that the functional impact of some QTL on septal traits will involve the collective effects of multiple variants on gene function and/or expression. This would represent a significant departure from a reductionist model whereby a single variant accounts for each QTL.

Our results provide the first high-resolution picture of genetic complexity and multilayered network liabilities underpinning atrial septal variation in a mouse model. It is noteworthy that 44% of high impact and predicted deleterious non-synonymous protein coding variants discovered in our model overlap with curated lists of known/high confidence and emerging human CHD gene lists. Thus, our study has strong relevance to the polygenic causation of CHD. Our results are consistent with Fisher’s ‘infinitesimal’ model whereby many loci could each contribute a small part to the liability for common disease (Fisher, 1918), and also GWAS analysis of human disease that most often detect common risk loci of small effect. However, as detected here and in our previous study (Kirk et al., 2006), individual septal QTL can contribute significant effects and allelic interactions are also likely important. Therefore, QTL studies in animal models have the potential to significantly inform our understanding of the genetic landscape underpinning human disease. Our quantitative model offers opportunities to study the interface between genetic, environmental, and epigenetic inputs to common CHD in greater detail.

## Materials and Methods

### QTL fine mapping using advanced intercross line

#### Mice and advanced intercross line

129T2/SvEms mice (substrain 129T2/SvEms-+^Ter^?) were obtained from The Jackson Laboratory and were a derivative of the Ter subline family of 129 strains (Biben et al., 2000). QSi5 is an inbred mouse strain that was bred from outbred Quackenbush Swiss mice (Holt et al., 2004) at the University of Sydney, Sydney NSW and were maintained as a breeding colony once established. QSi5 mice are also available from the Jackson Laboratory (strain 027001). As parental strains in our F2 study (Kirk et al., 2006), 129T2/SvEms and QSi5, they were selected based on having extreme values for mean FVL and prevalence of PFO, two indicators of atrial septal morphology which are strongly negatively correlated (Biben et al., 2000). 129T2/SvEms had the highest prevalence of PFO (75%) and shortest FVL (mean = 0.6 mm) among inbred strains and multiple crosses (Biben et al., 2000). QSi5 had the longest FVL (mean = 1.13 mm) and among the lowest prevalence of PFO (4.5%).

We randomly intercrossed the original F2 mice for 12 further generations. The F2 mice were bred to produce 48 male and female pairs which were then stocked in 48 separate cages. We intercrossed the mice in each cage to generate F3 mice. For the F3 × F3 and subsequent crosses, a cascading scheme was used in which a female mouse from one cage would be mated with a male mouse from the next cage. This breeding design, in which each pair contributes exactly two offspring (one male and one female) to the next generation doubles the effective population size and reduces the random changes in allele frequency due to random genetic drift over the 10 generations. Animals were bred and housed under Animal Care and Research Ethics approvals N00/4-2003/1/3745, N00/4-2003/2/3745 and N00/4-2003/3/3745 from the University of Sydney.

#### Dissection and measurements

In total, 1003 AIL F14 mice were dissected. Of these, 933 had complete phenotypic data (475 males and 458 females). Phenotyping of each mouse including initial and fine dissections, determination of PFO status and measurement of septal features, were all performed on the same day. As in the F2 study (Kirk et al., 2006), the thoracic organs including heart, lungs, and mediastinum were initially dissected *en bloc* and stored in PBS. A tail biopsy was also taken from each mouse and snap-frozen for DNA extraction.

The following steps were performed under a Leica MZ8 dissecting microscope. The mediastinal organs were removed to expose the atria. Subsequently, the left atrium was opened to expose the atrial septum. We detected PFO by pressurization of the right atrium. The right to left passage of blood across the inter-atrial septum indicated the presence of PFO. Measurement of septal features including FVL, FOW, and CRW was performed using an eyepiece graticule. FVL was defined formally, as in our previous study (Kirk et al., 2006), as the length of the flap valve from the edge of the crescent (proximal rim of the *ostium secundum*) to the distal rim of the fossa ovalis. The maximum width of the *foramen ovale* (*foramen ovale* width; FOW) was measured perpendicular to the FVL. Crescent width (CRW) was defined as the maximum width of the prominent crescent-shaped ridge, representing the proximal rim of the *ostium secundum* and edge of the flap valve as previously described (Kirk et al., 2006).

#### Normalization

A general linear model (PASW Statistics 18) was used to analyse the effect of various covariates on the traits of interest (FVL, FOW, and heart weight; HW) considering a *p*-value of < 0.05 as significant (**Supplementary Files 8-12**). FVL and FOW were significantly affected only by HW (*p* = 0.002 and *p* = 0.028, respectively) and the effect of other covariates including sex, age, body weight (BW), and coat colour was not significant. However, we did not adjust for HW since QTL relevant to HW may also influence atrial septal morphology. On the other hand, HW was significantly affected by sex (*p* < 0.001), age (*p* = 0.032), BW (*p* < 0.001) and coat colour (*p* = 0.042). Therefore, prior to sample selection and further analysis, HW was normalized for age, sex and BW, although not for coat colour so as to avoid missing QTL linked to coat colour genes.

#### Sample selection

Selective genotyping of extreme phenotypes is an efficient method to increase the power of QTL mapping (Lander and Botstein, 1989). However, the benefits of this method decline with an increasing number of uncorrelated or weakly correlated traits. The focus of our study was to fine map QTL underlying the main highly correlated traits (FVL and FOW) and a peripheral trait of HW. We did not consider CRW as a basis for sample selection since it was a less defined anatomical structure and, unlike FVL and FOW, it was not associated with PFO (see Results). For each trait (FVL, FOW, and HW), we selected approximately 100 F14 animals with extreme phenotype. Given the overlap between the extreme phenotypes from different traits, 237 mice were selected by this method. To compensate for biases in selective genotyping in this study, selection of extreme phenotypes was combined with a degree of random selection. Therefore, 163 mice were also selected randomly giving a total sample of 400 mice. The same number of males and females were selected from the breeding cages. Thus, the selected mice gave as equal as possible representation of males, females and breeding cages.

#### Marker selection

We searched genotype data from the Mouse HapMap project for potential informative single nucleotide polymorphisms (SNPs) between 129T2/SvEms and QSi5 strains. In total, a set of 135 markers with the average interval of 2 centimorgans (cM) was selected to genotype genomic regions including the significant QTL from the F2 study (three QTL for FVL and three for FOW) and one QTL for HW (**Supplementary File 13**). In addition, a suggestive QTL for FOW on MMU9 (LOD = 3.43) was included in the AIL study as its peak covered the known ASD/VSD gene *Tbx20* (Kirk et al., 2006). Fifteen extra markers were chosen to cover the whole peak of this QTL. Subsequently, a total of 150 markers were genotyped in the selected mice (**Supplementary File 13**).

#### Genotyping

Genomic DNA was isolated from mouse tails using Macherey-Nagel Nucleospin kit according to the manufacturer’s guidelines. Prior to genotyping, we verified the informativeness of the markers for parental strains. Subsequently, the markers were genotyped in the selected mice using iPLEX™ MassArray® assay following the manufacturer’s protocol.

#### Linkage analysis

We performed interval mapping linkage analysis for the quantitative traits at 0.25 cM intervals using a maximum likelihood method implemented in R software (see code availability statement below). The method used was initially developed for an F2 design, but modified for an AIL using the methods described by Darvasi and Soller (Darvasi and Soller, 1995). Given the density of selected markers and the size of the genomic region covered by markers a LOD score of 2 was set as the threshold level of significance (Lander and Botstein, 1989).

We used the 1-LOD drop-off to estimate the confidence interval of each QTL (Lander and Botstein, 1989). However, in some cases two significant peaks were located close together and overlapped in the 1-LOD drop-off intervals. To determine whether these peaks reflected the same underlying QTL or identified the presence of independent QTL, we re-ran the linkage analysis using a model in which the marker closest to the higher peak was included as a fixed term (**Figure 2-figure supplement 1**), effectively a simplified form of composite interval mapping but applied in a maximum likelihood framework (Kearsey and Hyne, 1994; Wu and Li, 1994). Disappearance of the lower peak would indicate that two peaks represented a single QTL. On the other hand, if the lower peak remained significant, two peaks would represent two separate QTL.

We have previously developed a QTL program in R to perform linkage analysis for PFO as a binary trait (presence or absence) (Moradi Marjaneh et al., 2012). For binary analysis of AIL data, the linear model used for the quantitative analysis was replaced by a generalized linear model in the form of a logistic regression model, as described previously (Moradi Marjaneh et al., 2012).

Additive and dominance effects given in **Supplementary File 3** were used to calculate QTL effect size information. The additive effect for each QTL (a.qtl) was defined as the effect of one copy of the Q (QSi5) allele and calculated as (mean.QQ - mean.qq)/2 where mean.QQ is the phenotypic mean for genotype QQ and mean.qq is the phenotypic mean for genotype qq (129T2/SvEms allele). The dominance effect (d.qtl) for each QTL was defined as the difference between the mean.Qq (phenotypic mean for genotype Qq) and the midpoint between the qq and QQ means. Therefore, if d.qtl > 0, allele Q is (partially) dominant as Qq is moved towards QQ and if d.qtl < 0, allele q is (partially) dominant as Qq is moved towards qq. Attributable phenotypic variance for each QTL is the additive effect of the QTL, expressed as a percentage of the difference between parental means for that trait.

#### Conversion of genetic maps

We previously defined the genetic positions of the F2 markers using an older mouse genetic map developed by the Whitehead Institute and the Massachusetts Institute of Technology (MIT) (Dietrich et al., 1996). Thus, we converted the genetic position of F2 markers into those for the current Mouse Genome Informatics (MGI) map developed at the Jackson Laboratory (Bult et al., 2008) using mouse maps introduced by Cox et al (Cox et al., 2009). For inter-marker intervals a linear interpolation was used to convert old genetic positions to new ones.

### Whole genome sequencing of the AIL parental strains

#### Sequencing

Genomic DNA was extracted from tail biopsy specimens of QSi5 and 129T2/SvEms mice using the phenol-chloroform extraction method. Library preparation and sequencing were performed at the Kinghorn Centre for Clinical Genomics. 200ng of DNA was mechanically fragmented using LE220 Covaris (Covaris, Woburn, USA) to approximately 450 bp inserts followed by library preparation using the Seqlab TruSeq Nano DNA HT kit (20000903, Illumina, San Diego, USA). Library preparation was performed according to manufacturer’s instructions which included end repair, library size selection, 3′ end polyadenylation, adaptor ligation, purification of ligated fragments, enrichment of ligated fragments by PCR and a final purification of amplified DNA library. Library fragment sizes were reviewed using the LabChip GX (Perkin Elmer, Waltham, USA) to evaluate library size and adapter dimer presence before the library concentration was determined via quantitative real time PCR using the KAPA library quantification kits (KK4824, Roche, Basel, Switzerland) on the QuantStudio7 or ViiA7 (Thermo Fisher Scientific, Waltham, USA). Normalised DNA libraries were clustered on Illumina cBot then sequenced using Illumina HiSeq X Ten platform using HiSeq X Ten Reagent Kit v2.5 kits (FC-501-2501, Illumina). Paired end sequencing was performed using the 2×150bp chemistry to achieve an average output of approximately >120 Gb of data per library.

#### Bioinformatic analysis

The sequencing reads were mapped to the mouse reference genome (NCBI38/mm10) using Burrows-Wheeler Aligner (BWA-mem v0.7.15) (Li and Durbin, 2010). Variants were called using the genome analysis toolkit (GATK; v3.5) pipeline following best practice guidelines (van der Auwera and O’Connor, 2020). This involved marking duplicate reads with the Picard toolkit (Broad Institute GitHub Repository: http://broadinstitute.github.io/picard/), indel realignment with GATK using known SNPs and indels from C57BL/6 dbSNP142 (ftp://ftp-mouse.sanger.ac.uk/current_indels/strain_specific_vcfs/), Base quality score recalibration and final variant calling was peformed using the GATK Haplotype Caller. Variant call format (VCF) files were merged using VCFtools (v0.1.14) (Danecek et al., 2011) and annotated with ANNOVAR (Wang et al., 2010). A set of high quality variants was identified by filtering variants for those with a minimum sequencing depth of 8 and maximum estimated false discovery rate (FDR) of 10%. Variants were filtered to only consider those that were homozygous in one mouse line with either the reference or a heterozygous call in the other line. Zygosity information from each mouse line was extracted using VCFtools and custom scripts.

To analyse protein-coding variants, variants were filtered for feature annotations ‘exonic’, ‘exonic;splicing’ or ‘splicing’, with variants annotated with effect class ‘synonymous SNV’ filtered out. Variants were filtered for genes expressed in the septal time course RNA-seq (see below), where a gene was defined as expressed if it had a counts-per million (CPM) > 1 in at least 2 samples. They were then converted to mm10/GRCm39 coordinates and submitted to the Ensembl Variant Effect Predictor (VEP) tool (McLaren et al., 2016). Based on VEP annotations, variants were classified as high-impact (VEP impact classification of ‘HIGH’), deleterious missense (SIFT (Ng and Henikoff, 2003) ‘deleterious’ prediction) or non-high-impact deletions. High-impact variants were manually inspected in IGV and filtered out if the ‘HIGH’ impact classification was deemed inaccurate (for example, cases where an adjacent variant negated a predicted stop gain or a substitution was incorrectly called as a deletion). Finally, the list of high-confidence potential-pathogenic variants were filtered for variants falling within QTL coordinates (LOD > 1).

### Transcriptome analysis of cardiac interatrial septum in mice

Microdissection of the atrial septum region was performed on embryos from mouse strains (QSi5, 129T2/SvEms) at E12.5, E14.5 and E16.5 (six septa per mouse strain per time point). Embryos were dissected from the uterus in ice-cold PBS under a dissecting scope (Leica MZ8) using micro-tweezers. Once the heart was harvested from the embryo and pericardial membrane removed, the atrial and ventricular regions were separated. Then, the atrial appendages were removed from the atrial part and remaining mesenchymal tissue from the atrioventricular canal further trimmed. The remaining tissue, containing the atrial septum, was stored at −80°C in RNAlater solution (Ambion). Total RNA was purified using miRNeasy Micro Kit (Qiagen). Two technical replicates per mouse strain per time point were created by pooling RNA from three dissected septa each.

#### RNA sequencing and data analysis

RNA libraries were made using TruSeq stranded RNA Library Preparation Kit (Illumina) and sequenced on a HiSeq2500 Illumina sequencer to a depth of ∼23 million paired end reads per sample. Sequencing reads were trimmed to remove poor quality sequence and adaptors using Trimmomatic (v0.35) (Bolger et al., 2014) using parameters LEADING:3 TRAILING:3 SLIDINGWINDOW:4:15 MINLEN:33.). Sequencing reads were aligned against the mouse reference genome (GRCm38/mm10) using RNA STAR (v2.5.1) (Dobin et al., 2013). Gene counts were assembled using SummariseOverlaps (v1.30.0) (Lawrence et al., 2013) against Gencode release M4 and filtering to only include those whose expression was at least 1 count per-million (CPM) in at least two samples. Differential expression analysis of RNA-seq data was performed using DESeq2 (v. 1.14.1) (Love et al., 2014). For differential expression analysis, each time-point was analysed individually. Differentially expressed genes were calculated between 129T2/SvEms and QSi5 using the *DESeq* function with default parameters, with an adjusted *p*-value cut-off of 0.05 and absolute log2 fold-change difference of 0.5 used as thresholds for significance. GO term analyses of differentially expressed genes was performed using the PANTHER web-service (Mi et al., 2017) with a false discovery rate cut-off of 0.05 used to assess significance.

#### Network analysis

Network analysis of DEGs was performed using protein-protein association connections obtained from the STRING (Szklarczyk et al., 2017) version 10 database, considering connections with a combined score greater than 700. Permutation testing (100,000 permutations) was applied to determine whether the size and complexity of the networks (determined by number of edges and clustering coefficient, respectively) were greater than expected by chance. For each permutation, a random set of genes was selected from the set of expressed genes in the RNA-seq for the relevant time-point, and network metrics re-calculated. Edge counts and clustering coefficients were calculated using NetworkX (Aric A. Hagberg, 2008). Empirical *p*-values were then determined as the proportion of random networks achieving network metrics equal to, or higher, than the DEG networks.

### Analysis of SMAD6 variant

#### Plasmids

Pcs2-SMAD6 (#14960) and BRE-luc (#45126) were purchased from Addgene. FLAG tag was added to SMAD6 using following primers: Forward (F): 5’ GATCGACTACAAGGACGACGATGACAAGG 3’; Reverse (R): 5’ GATCCCTTGTCATCGTCGTCCTTGTAGTC 3’.

#### Mutagenesis primers

R281P - F 5’ CTCCCTACTCTCCGCTGTCTCCTCG 3’; R 5’ CGAGGAGACAGCGGAGAGTAGGGAG 3’; Y279A - F 5’ CTCCGCCACCTCCCGCATCTCGGCTGTCTC 3’; R 5’ GAGACAGCCGAGATGCGGGAGGTGGCGGAG 3’; G471S - F 5’ CATCAGCTTCGCCAAGAGCTGGGGGCCCTG 3’; R 5’ CAGGGCCCCCAGCTCTTGGCGAAGCTGATG 3’.

#### Cell culture

HEK293T cells (ATCC; www.atcc.org) were maintained in DMEM medium containing 10% FCS in a humidified incubator at 37°C at 10% CO2. 80,000 cells were seeded in 12-well plates for luciferase assays and 100,000 cells were seeded in 6-well plates for protein extraction. BMP4 ligand (Gibco, #PHC9534) was added to cells at 100 ng/ml. Transfections were carried out using Lipofectamine 3000 (ThermoFisher Scientific) according to manufacturer’s instructions.

#### Luciferase assay

HEK293T cells were transfected with 100 ng of luciferase reporter constructs, 150 ng of expression vectors or empty vector, and 2.5 ng of the TK-Renilla. Cells were treated with BMP4 overnight on the day of transfection. Assays were performed 24 hrs after transfection. Dual luciferase assays were performed as per manufacturer’s instructions (Promega, #E1980). Firefly luciferase activity was normalized to Renilla luciferase activity.

#### Co-IP and western blot

HEK293T cells were transfected with 1 µg of hSMAD6-WT, hSMAD6-R281P, hSMAD6-Y279A, or hSMAD6-G471S. Cells were treated with either DMSO or MG132 (Sigma Aldrich, #M7449) 8 hrs prior to protein extraction. Cells were lysed 48 hrs after transfection using whole cell extract buffer (20 mM HEPES, 420 mM NaCl, 0.5% NP-40, 25% glycerol, 0.2 mM EDTA, 1.5 mM MgCl2, 1 mM PMSF and protease inhibitors). Cells were washed in PBS and lysed with 250 µl WCE buffer for 10 mins on ice. Lysed cells were scraped off the 6-well plates and homogenized with a 25 Gauge needle 10 times. The lysates were centrifuged for 30 minutes at 4°C. Supernatant was precleared with Protein G (Thermo Fisher Scientific) for 1 hr at 4°C, then incubated with anti-FLAG M2 antibody (Sigma Aldrich, #F1804) at 1:150 dilution or an equal amount of mouse IgG overnight at 4°C. The lysates were incubated with Protein G beads for 2 hrs at 4°C, then washed in WCE buffer 4 times. Protein was eluted in 4x sample buffer (Biorad) for 5 mins at 95°C and loaded onto TGX stain-free precast gels (Biorad). Western blots were carried out using the following antibodies: anti-ubiquitin (1:1000, Santa Cruz Biotechnology, sc-166553), anti-FLAG (1:1000, Cell Signaling Technology, #14793), anti-SMAD6 (1:100, Santa Cruz Biotechnology, sc-25321).

#### Variant enrichment analyses

We examined the overlap of high quality homozygous 129T2/SvEms vs QSi5 variants and genomic features (enhancers, promoters, 5’UTRs, exons, introns, and 3’UTRs) of DEGs and non-DEGs located under QTL 1-LOD drop-off regions. Promoters were defined as 1000 bp upstream and 100 bp downstream of a known transcription start site and enhancers were defined as H3K4me1^+^ or H3K27ac^+^, and H3K4me3^-^ regions 50 kb upstream of transcription factor start sites. Accession numbers for histone mark data were: H3K4me1 - ENCFF817VKF; H3K27ac - ENCFF815RLC); HK4me3 – ENCFF994HEQ. Variant enrichment within each feature was defined as the proportion of variants that overlapped the feature, with normalisation for the proportion of coverage of the feature within the QTL 1-LOD drop-off. To test the significance of the enrichment, background controls were generated by permuting each genomic feature of DEGs and non-DEGs under QTL, respectively, by randomly selecting matched size blocks (according to the tested property) under QTL regions. This process was repeated 1000 times using *bedtools shuffle* (version 2.25.0). For each genomic feature, the DEG/non-DEG regions were excluded from the pool of regions used to generate random interval sets. The overlap of the variants and genomic features of DEGs and non-DEGs under QTL was compared to the mean overlap for 1000 random interval sets and empirical *p*-values were calculated by dividing the number of random interval sets showing equal or greater overlap than the observed by 1000.

## Data availability

Sequencing data have been deposited in the ArrayExpress database at EMBL-EBI (www.ebi.ac.uk/arrayexpress) under accession codes E-MTAB-11161 (DNA-seq) and E-MTAB-10929 (RNA-seq).

## Code availability

The codes used to perform linkage analysis are available at https://github.com/MahdiMoradiMarjaneh/AIL

## Acknowledgments

MMM held a University of New South Wales (UNSW Sydney) International Postgraduate Award (ID3263695). EPK held a National Heart Foundation Clinical Fellowship. The work was supported by the National Institute of Heart Lung and Blood (USA; 1RO1HL68885-01), National Heart Foundation of Australia (G06S2575; G0050738), National Health and Medical Research Council (NHMRC, Australia) (354400, 0573732; 1074386), the New South Wales Government Ministry of Health 20:20 campaign and the Victor Chang Cardiac Research Institute Innovation Centre (funded by the New South Wales Government Ministry of Health). RPH was supported by an NHMRC Australia Fellowship (0573705) and Senior Principal Research Fellowship (1118576). PCP was supported by a UNSW Sydney International Postgraduate Award (UIPA) PhD Scholarship.

## Supplementary Figure Legends

**Figure 1-figure supplement 1.**
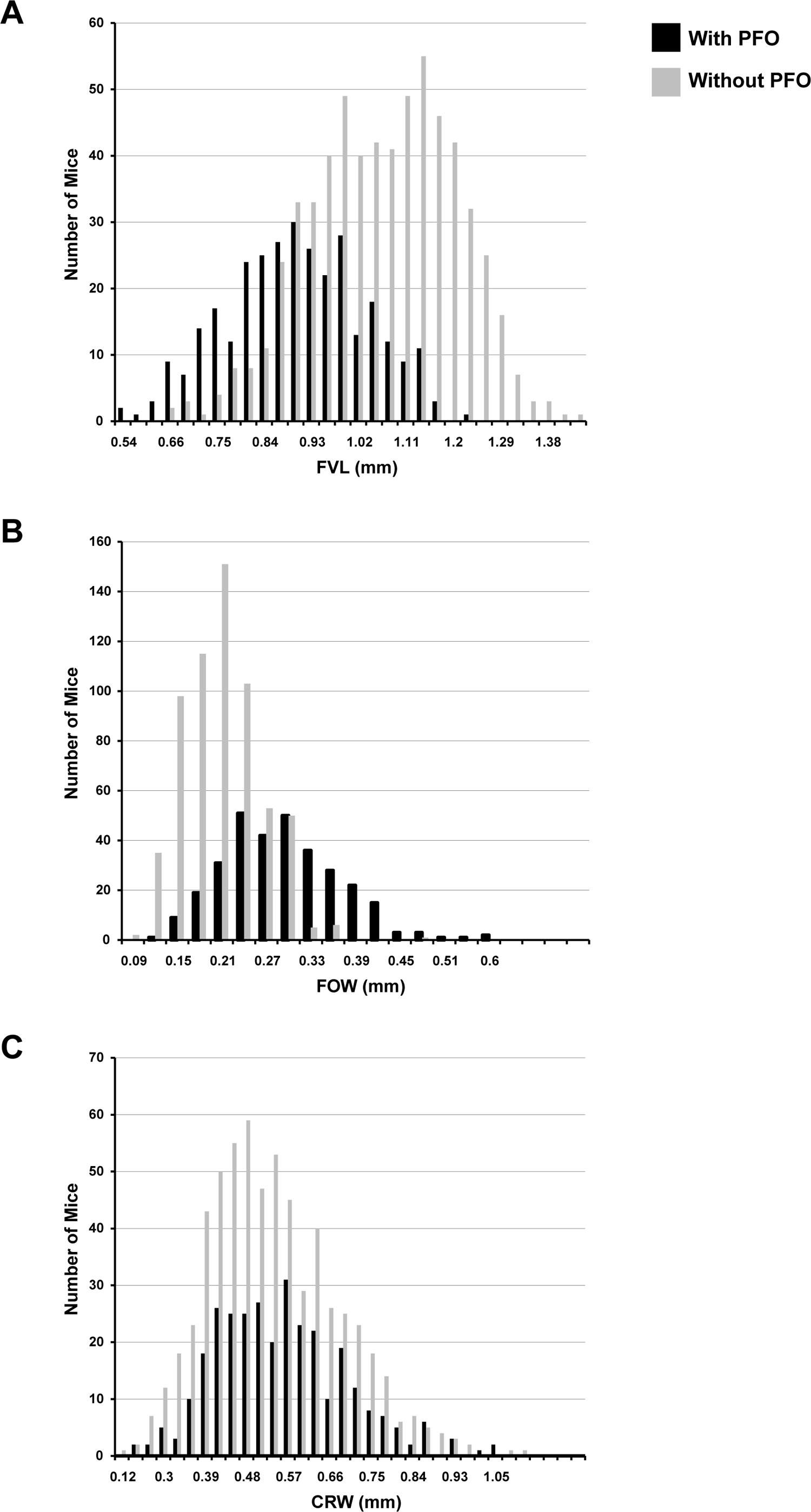
Histograms of quantitative traits in F14 mice with and without PFO.

**Figure 2-figure supplement 1.**
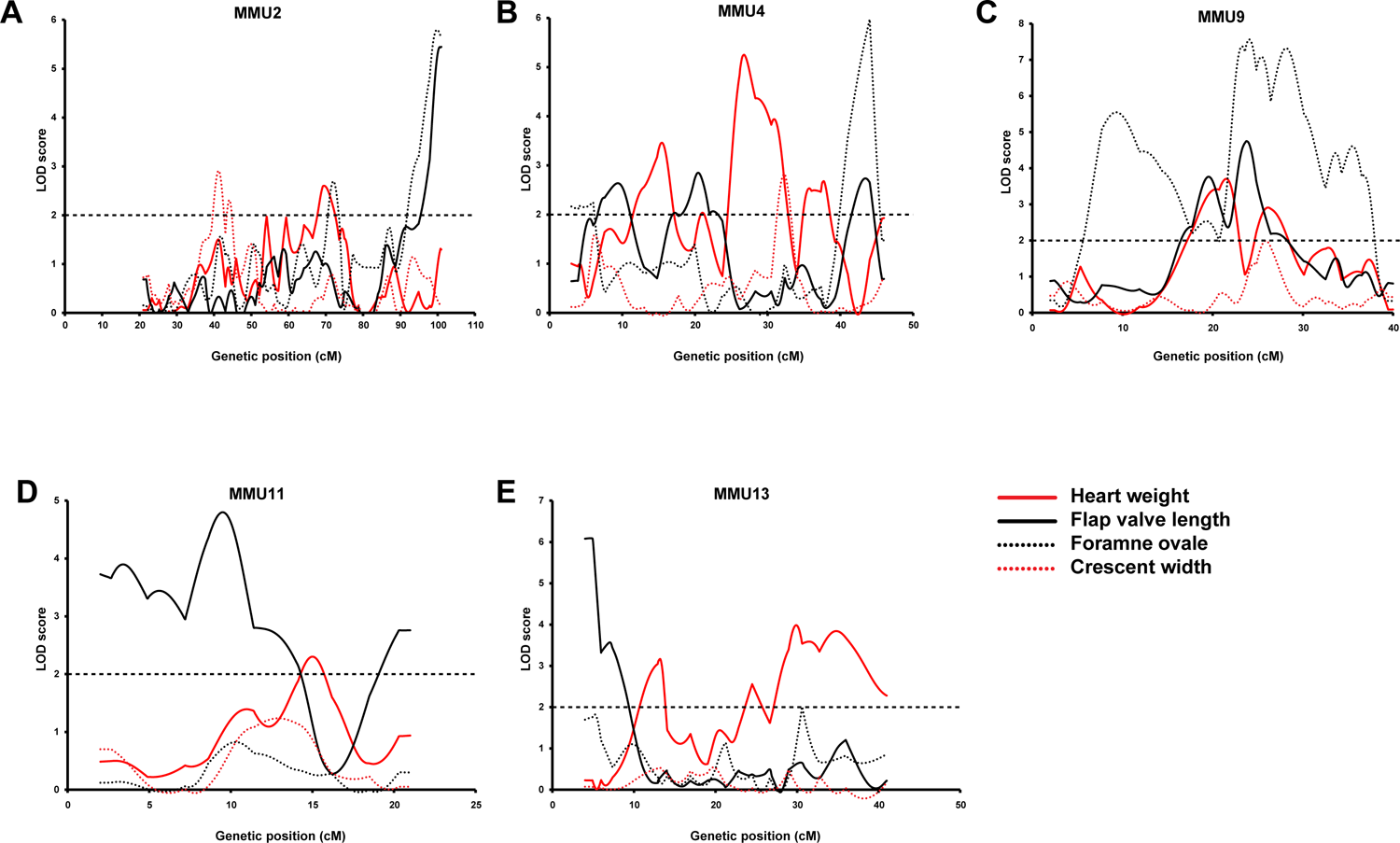
Comparison of AIL results for HW (red line) and quantitative traits of atrial septum.

**Figure 3-figure supplement 1.**
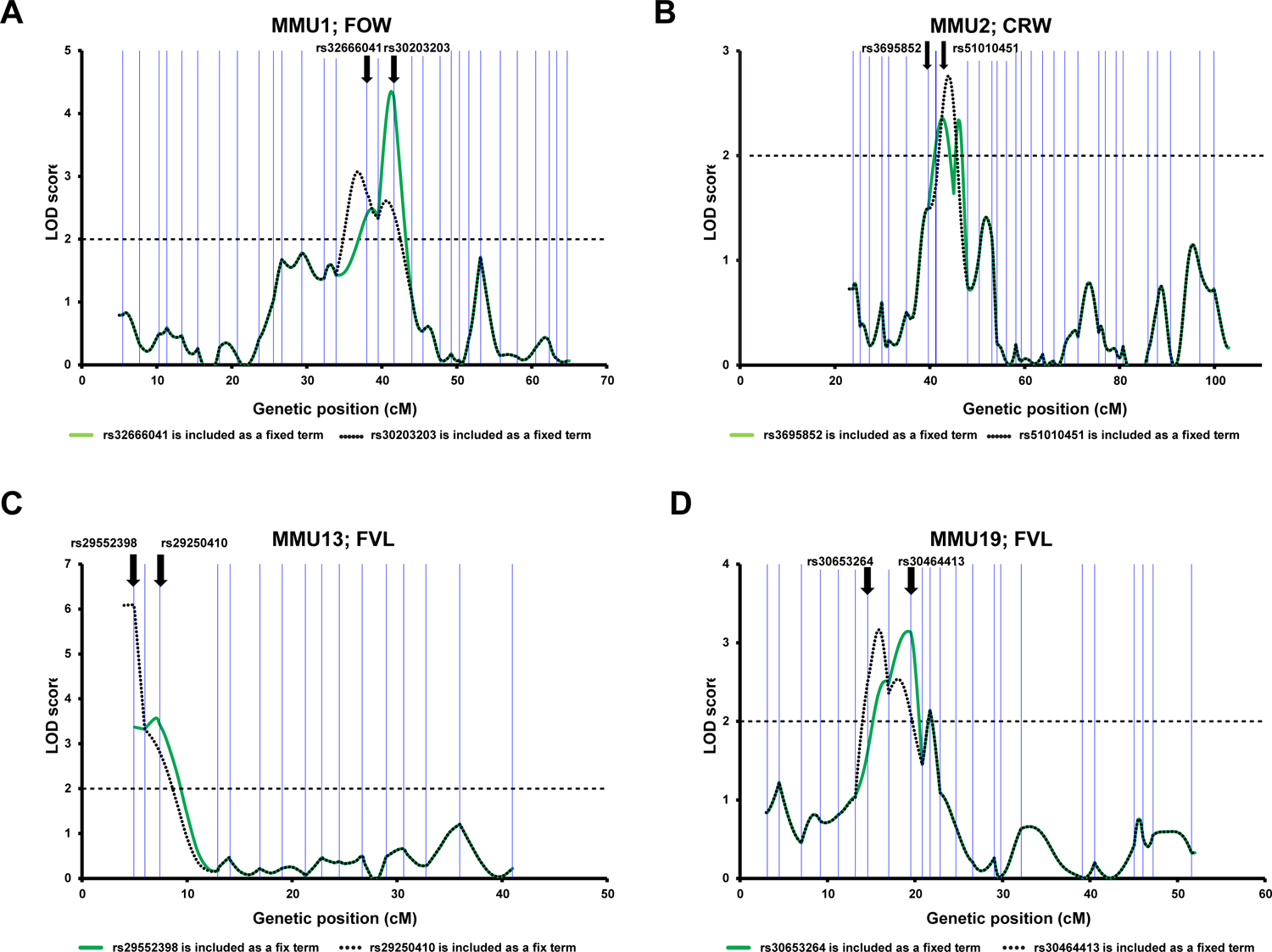
Linkage analysis of the AIL data. For each chromosome two different models have been used in each, one peak marker was included as a fixed term. Vertical blue lines correspond to the positions of the AIL markers. The bold arrows indicate the positions of the fixed markers.

**Figure 5-figure supplement 1.**
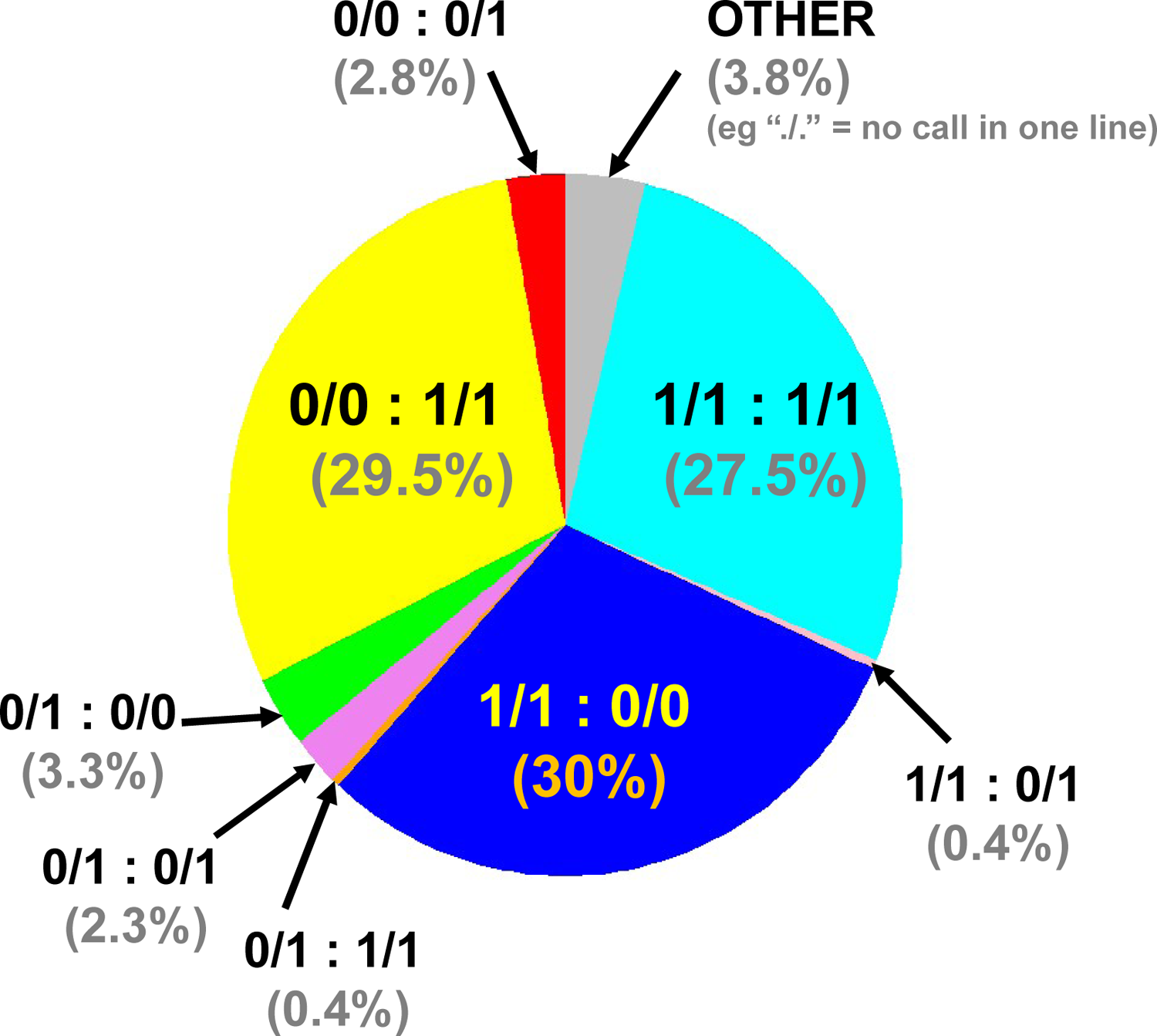
Combined genome-wide variant zygosity of AIL parental strains. The majority of SNPs are either unique to 129T (1/1:0/0; 30%; dark blue) or QSi5 (0/0:1/1; 29.5%; yellow) or common to both lines (1/1:1/1; 27.5%; light blue) with respect to the C56Bl/6 reference. “Other” includes, for example ./. = one call in one line. The low number of heterozygous calls may be annotation artifacts due to the presence of repetitive sequences.

**Figure 5-figure supplement 2.**
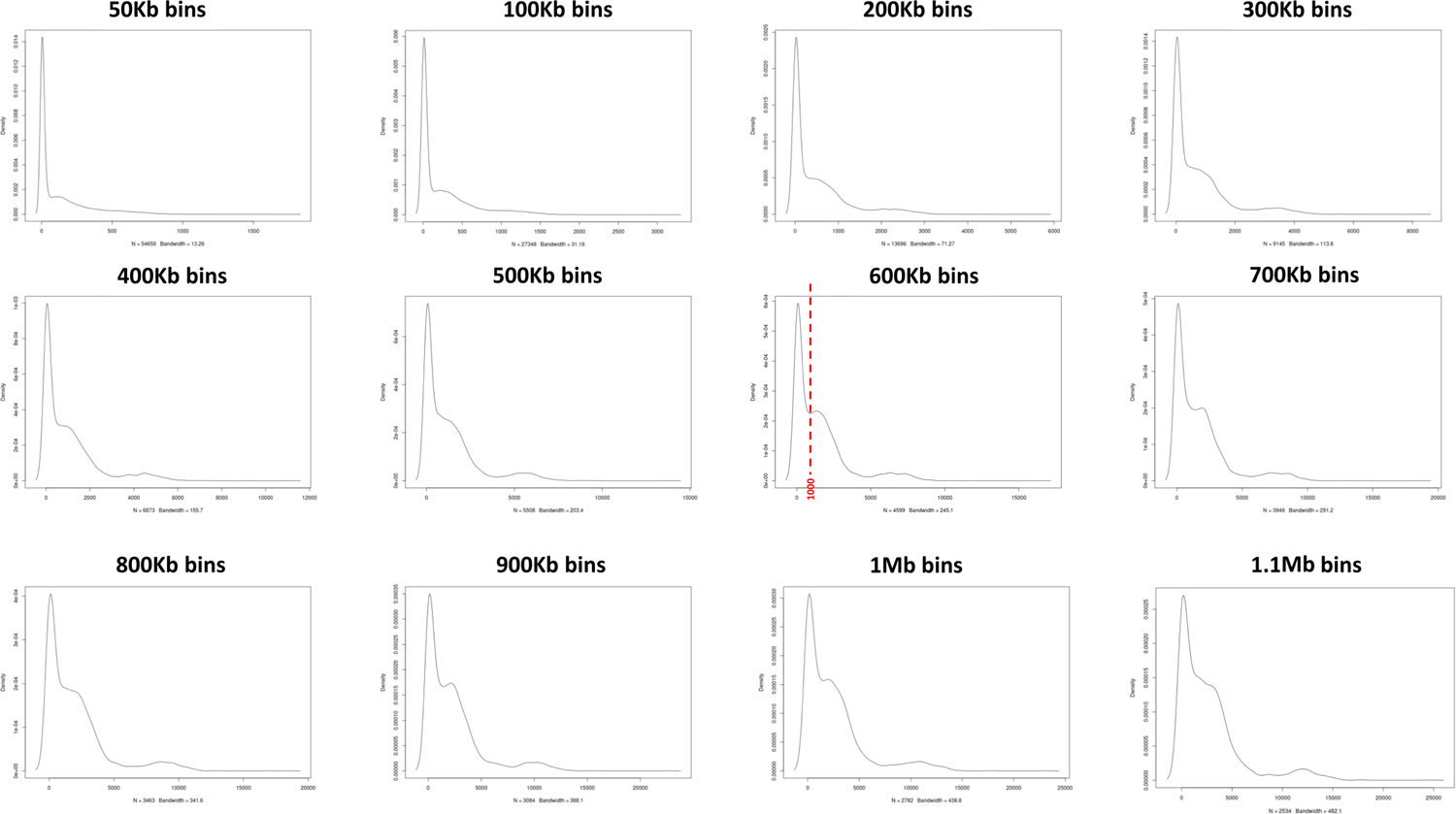
Assessment of 129T2/SvEms vs QSi5 variant density across the genome using different bin sizes. A bin of 600kb provided the clearest distinction between high and low variant density peaks defining a cutoff of 1000 variants/600Kb.

**Figure 8-figure supplement 1.**
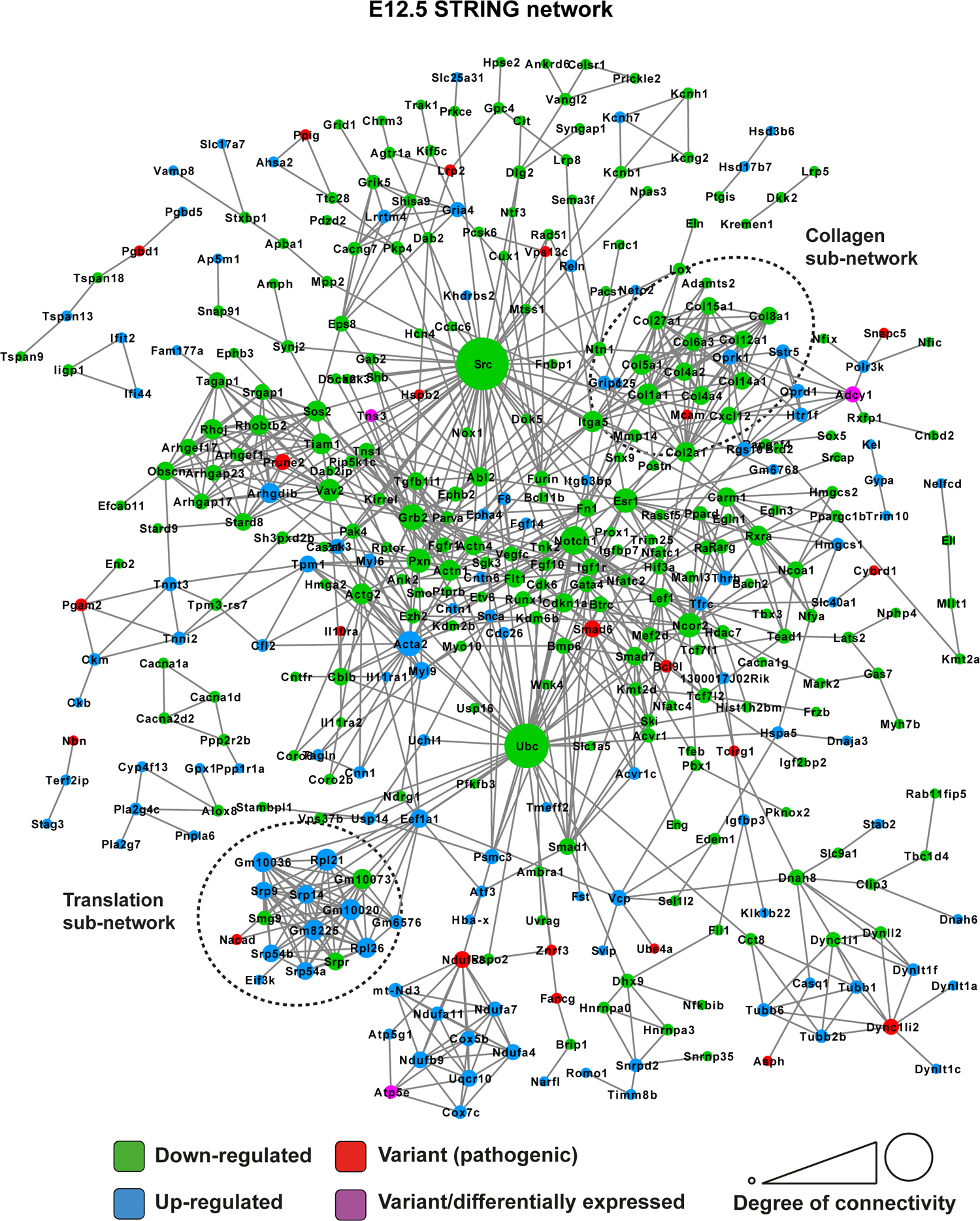
STRING network of E12.5 differentially expressed genes and genes with predicted-pathogenic variants from QSi5 or 129T2/SvEms. Genes are coloured according to whether they are upregulated or downregulated in 129T2/SvEms, contain a predicted-pathogenic variant, or are both differentially expressed and containing a variant.

**Figure 8-figure supplement 2.**
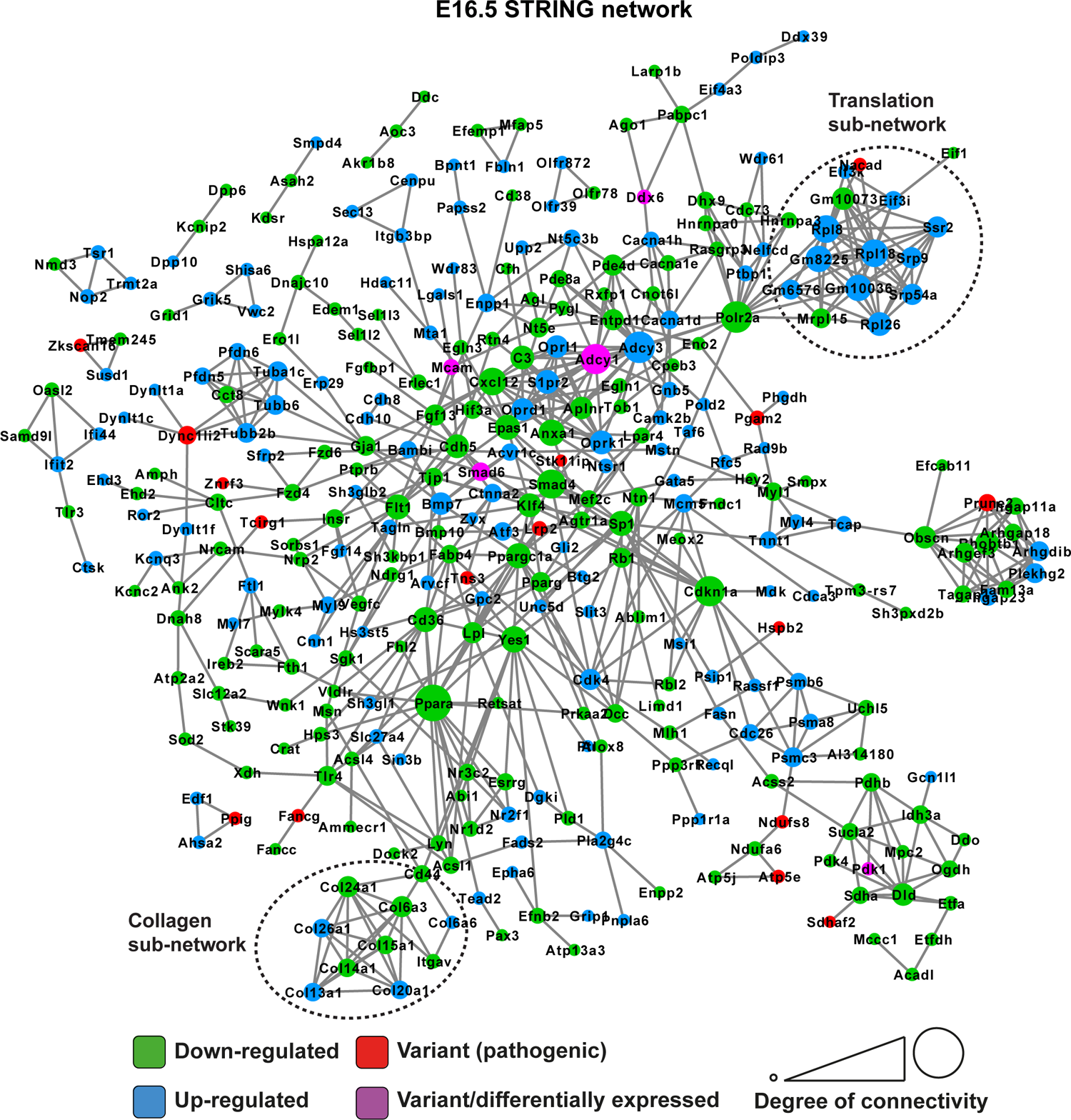
STRING network of E16.5 differentially expressed genes and genes with predicted-pathogenic variants from QSi5 or 129T2/SvEms. Genes are coloured according to whether they are upregulated or downregulated in 129T2/SvEms, contain a predicted-pathogenic variant, or are both differentially expressed and containing a variant.

**Figure 10-figure supplement 1.**
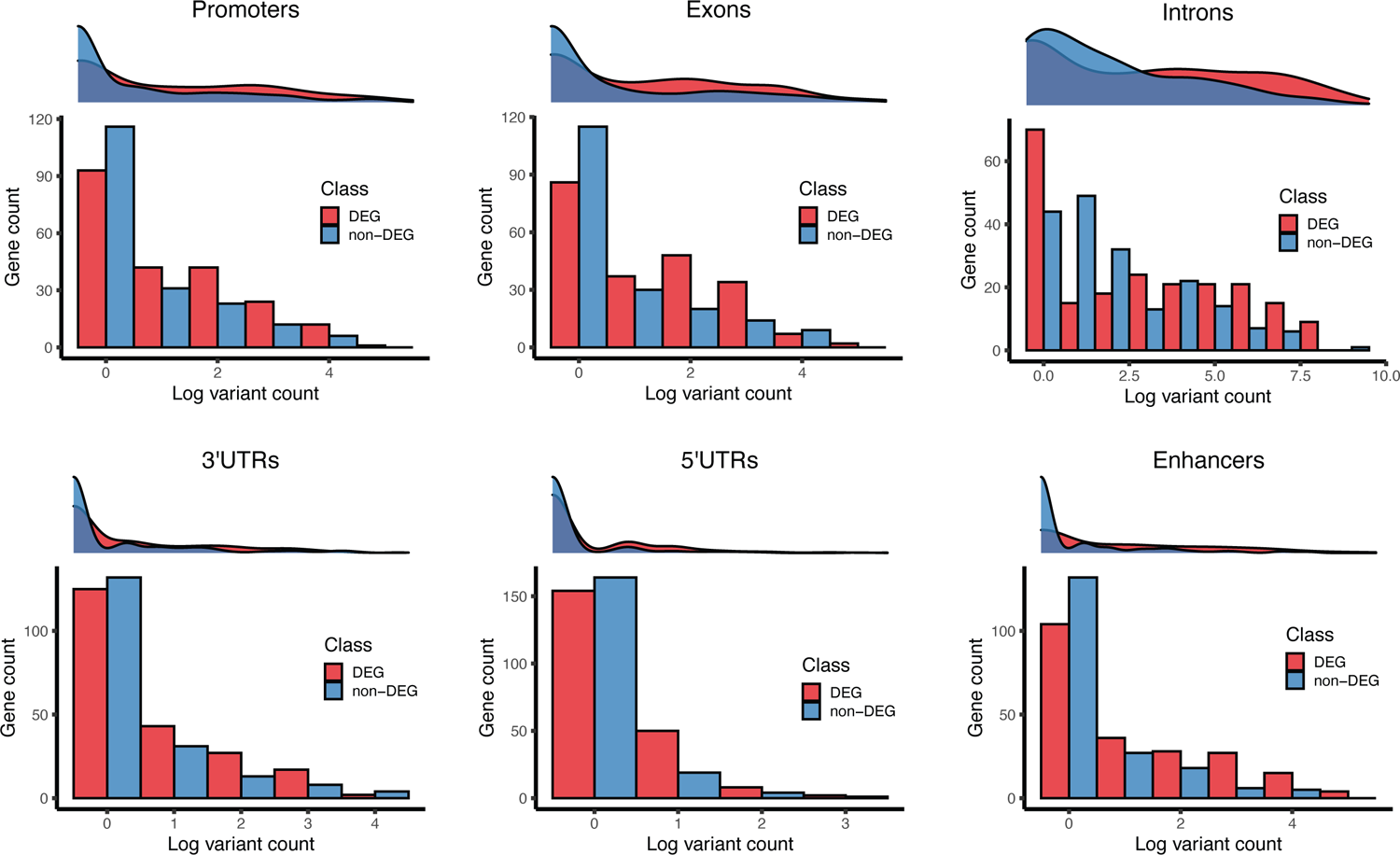
Gene counts by number of 129T2/SvEms vs QSi5 variants. For each genomic feature, a density plot and histogram representing the log-transformed number of variants per gene (x-axis) and frequency of the genes (y-axis). Each plot illustrates the gene counts for DEG (red) and non-DEG (blue) genes under QTL.

## Figure Source File Legend

**Figure 7-source data 1.** Original blots associated with Figure 7.

## Supplementary File Legends

**Supplementary File 1.** Relationship between PFO and the quantitative traits in F14 mice with complete data (n = 933).

**Supplementary File 2.** Inter-trait correlation coefficients (r) in F14 mice with P-values in brackets.

**Supplementary File 3.** QTL identified by AIL study.

**Supplementary File 4.** High impact, deleterious missense, and deletion variants between 129T2/SvEms and QSi5 strains.

**Supplementary File 5.** Differentially expressed genes (DEGs) between 129T2 vs QSi5 mice across the developmental time points.

**Supplementary File 6.** Gene ontology biological process terms that were significantly over-represented among the DEGs.

**Supplementary File 7.** Counts of variants in different genomic features for DEGs and non-DEGs.

**Supplementary File 8.** Effect of various covariates on FVL.

**Supplementary File 9.** Effect of various covariates on FOW.

**Supplementary File 10.** Effect of various covariates on CRW.

**Supplementary File 11.** Effect of various covariates on HW.

**Supplementary File 12.** Effect of various covariates on PFO.

**Supplementary File 13.** List of markers with physical and genetic location.

## Notes

### Competing Interest Statement

The authors have declared no competing interest.

